# Modelling cell-cell interactions from spatial molecular data with spatial variance component analysis

**DOI:** 10.1101/265256

**Authors:** Damien Arnol, Denis Schapiro, Bernd Bodenmiller, Julio Saez-Rodriguez, Oliver Stegle

## Abstract

Technological advances allow for assaying multiplexed spatially resolved RNA and protein expression profiling of individual cells, thereby capturing physiological tissue contexts of single cell variation. While methods for the high-throughput generation of spatial expression profiles are increasingly accessible, computational methods for studying the relevance of the spatial organization of tissues on cell-cell heterogeneity are only beginning to emerge. Here, we present *spatial variance component analysis (SVCA),* a computational framework for the analysis of spatial molecular data. SVCA enables quantifying the effect of cell-cell interactions, as well as environmental and intrinsic cell features on the expression levels of individual genes or proteins. In application to a breast cancer Imaging Mass Cytometry dataset, our model allows for robustly estimating spatial variance signatures, identifying cell-cell interactions as a major driver of expression heterogeneity. Finally, we apply SVCA to high-dimensional imaging-derived RNA data, where we identify molecular pathways that are linked to cell-cell interactions.

## Introduction

Experimental advances have enabled assaying RNA and protein abundances of single cells in spatial contexts, thereby allowing to study single cell variation in tissues. Already, these technologies have delivered new insights into tissue systems and the sources of transcriptional variation (Battich, Stoeger, & Pelkmans, 2013; Bodenmiller, 2016), with potential use as biomarkers for human health (Bodenmiller, 2016). Spatial expression variation can reflect interactions between adjacent cells, or can be caused by cells that migrate to specific locations in a tissue to perform their functions (e.g. immune cells).

Different technologies allow for generating spatially resolved expression profiles. Imaging Mass Cytometry (IMC) (Giesen et al., 2014) and Multiplexed Ion Beam Imaging (MIBI) (Angelo et al., 2014) rely on protein labeling with antibodies coupled with metal isotopes of specific masses followed by high-resolution tissue ablation and ionisation. IMC enables profiling of up to 27 targeted proteins with subcellular resolution. Other methods such as MxIF and CycIF use immunofluorescence for protein quantification of tens of markers at a time (Gerdes et al., 2013; Lin, Fallahi-Sichani, & Sorger, 2015). Increasingly, there also exist optical imaging-based assays to measure single cell RNA levels. Mer-FISH and seq-FISH use a combinatorial approach of fluorescence-labeled small RNA probes to identify and localise single RNA molecules (Chen, Boettiger, Moffitt, Wang, & Zhuang, 2015; Gerdes et al., 2013; Lin et al., 2015; Shah, Lubeck, Zhou, & Cai, 2017), which allows for a larger numbers of readouts (currently between 130 and 250). Even higher-dimensional expression profiles can be obtained from spatial expression profiling techniques such as Spatial Transcriptomics (Ståhl et al., 2016), which currently however do not offer single cell resolution and are therefore not adequate to study cell-to-cell variation.

The availability of spatially resolved expression data represents an unprecedented opportunity to disentangle largely unexplored sources of single cell variations: i) intrinsic sources of variation due to difference in cell types or states (e.g. cell-cycle state), ii) environmental effects from the cell microenvironment and iii) interactions between adjacent cells. Yet, although experimentally spatial omics profiles can be generated with high throughput, the required computational strategies for interpreting the resulting data are only beginning to emerge. On the one hand, there exist methods to cluster the cells based on their expression profiles (Achim et al., 2015), and there also exist statistical tests to assess the overall effect of the spatial topology on gene expression (Svensson, Teichmann, & Stegle, n.d.). However, these methods do not allow for directly assessing cell-cell interactions. On the other hand, there are methods that explain dependencies between cells based on cell-type assignments and predefined cellular neighbourhoods (Schapiro et al., 2017; Schulz et al., 2018). While these approaches provide qualitative insights into interactions between cell types, they do not allow for quantifying the impact of spatial effects on individual genes or proteins. Finally, there are regression-based models to assess interaction effects based on predefined features of cell neighbourhood (Battich, Stoeger, & Pelkmans, 2015; Goltsev et al., 2018), which however require discretization steps (see **Methods** for a detailed comparison).

Here, we present *Spatial variance component analysis (SVCA),* a computational framework to model spatial sources of variation of individual genes or proteins. SVCA allows for decomposing the sources of variation into intrinsic effects, environmental effects and cell-cell interactions. The model is parameterized based on continuous expression profiles of individual cells, and in particular avoids the need to define discrete cell types and microenvironmental variables. We illustrate SVCA using data from different technologies and biological domains, including IMC proteomics profiles data from human cancer tissue and spatial single-cell RNA profiles from the mouse hippocampus generated using seqFISH. Across these applications, we find that cell-cell interactions substantially contribute to gene expression variability, and we identify biologically relevant genes and pathways that participate in these interactions.

## Results and Discussion

SVCA considers random effect components that capture i) *intrinsic* sources of variation due to differences in cell types or states, ii) *environmental* sources of variation due to local extracellular factors iii) source of variations due to *cell-cell interactions* (**Fig. 1a**). The contribution of these factors on expression variability is modelled using additive random effects (**Fig. 1b**), where we borrow covariance models that were originally designed for social genetic effects (Baud et al., 2017) to capture cell-cell interactions. Importantly, SVCA does not require assigning cells to discrete types, but instead is based on a continuous measure of cell-cell similarities that are directly estimated from cell expression profiles (**Fig. S1**). The model also circumvents the need to define discrete neighbourhoods but instead weights interactions between pairs of cells as a function of their distance (**Fig. S1**). Once fitted, SVA allows for decomposing the sources of variation of individual genes and proteins (**Fig. 1c**) and the model can be used to assess the significance of these respective components (**Methods**).

**Figure 1.**
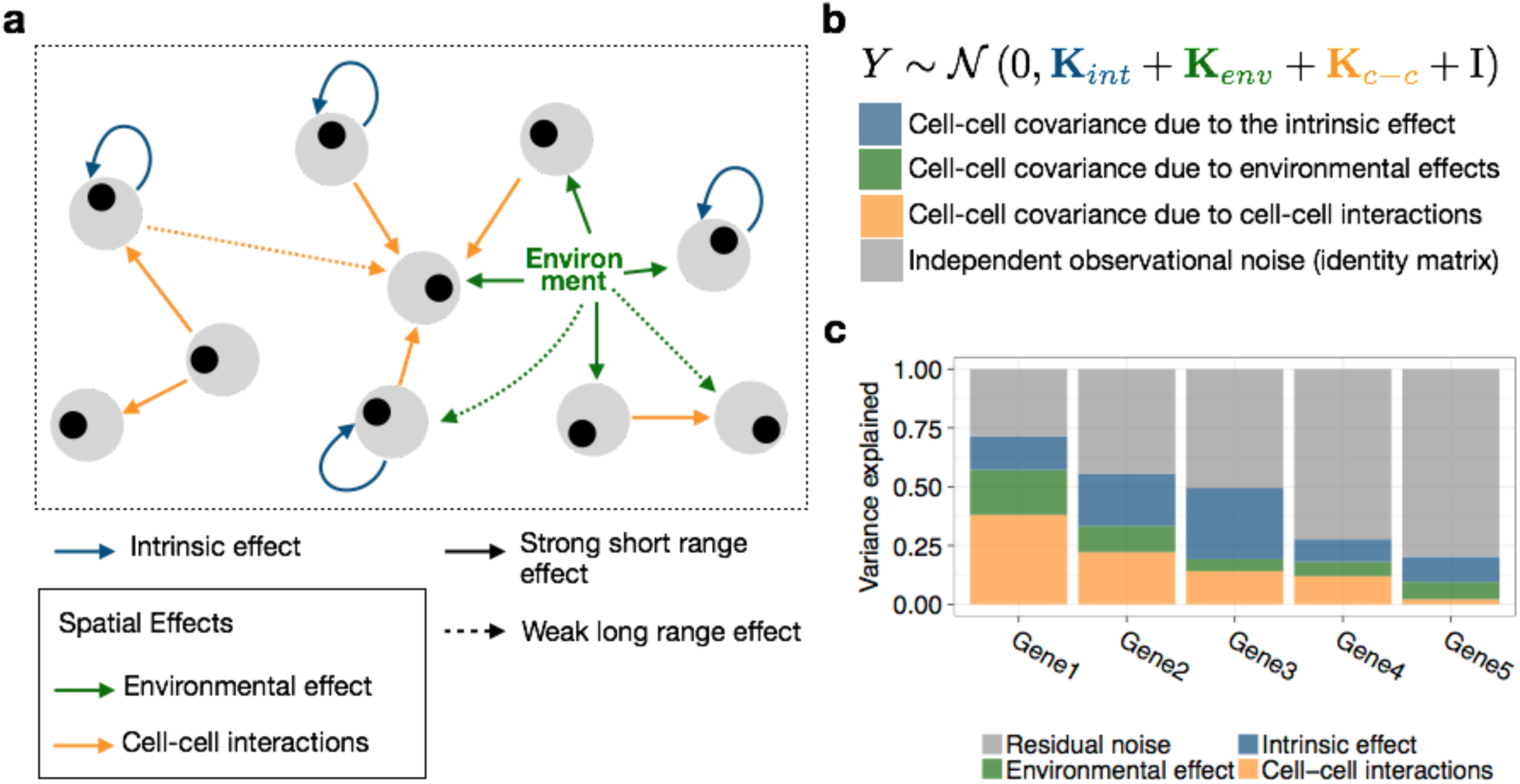
Spatial variance component analysis (SVCA): approach and overview. (**a**) SVCA decomposes the variability of individual genes and proteins into i) cell intrinsic effects (due to differences in intrinsic cell type or state, blue), ii) general environmental effects that capture expression differences due to local extracellular factors (green) and iii) a cell-cell interaction component that captures differences in expression level attributable to different cellular composition of a cell’s neighbourhood (yellow). (**b**) SVCA builds on a random effect framework to model additive contributions of these components. See **Fig. S1** and **Methods** for details on the definition of the corresponding random effects. (**c**) SVCA output: gene-level break down of the proportion of variance attributable to different components.

Initially, we used simulated data to assess the statistical calibration and the accuracy of SVCA variance estimates. Briefly, we generated expression profiles by sampling from the model, using empirical parameters derived from 11 real datasets (**Methods**), including the empirical position of cells, cell state covariance and ranges of fractions of variance explained by different components that reflect those observed in real data. First, we simulated expression profiles with no interaction effects to assess the calibration of the corresponding test, finding that the model yields conservative estimates (**Fig. 2a**). We also assessed the detection power for the test for cell-cell interactions, when varying the variance explained by this effect (**Fig. 2c**), and when varying the number of cells in the dataset (**Fig. 2d**). We also compared the estimated variance components for cell-cell interactions with the simulated variance components, again observing that the model estimates are conservative (**Fig. 2b**). To investigate the empirical identifiability of cell-cell interactions versus environmental effects, we also compared the estimates of the full model to a reduced model without the cell-cell interaction component. The results obtained show that the environmental effect harvests spatial sources of variation when they are not specifically modelled as in the cell-cell interaction term. This suggests that cell-cell interaction components are not confounded by other spatial sources of variation and are therefore conservative. Overall, we demonstrate that SVCA can be used to estimate and test for spatial drivers of single cell variability, in particular cell-cell interactions.

**Figure 2.**
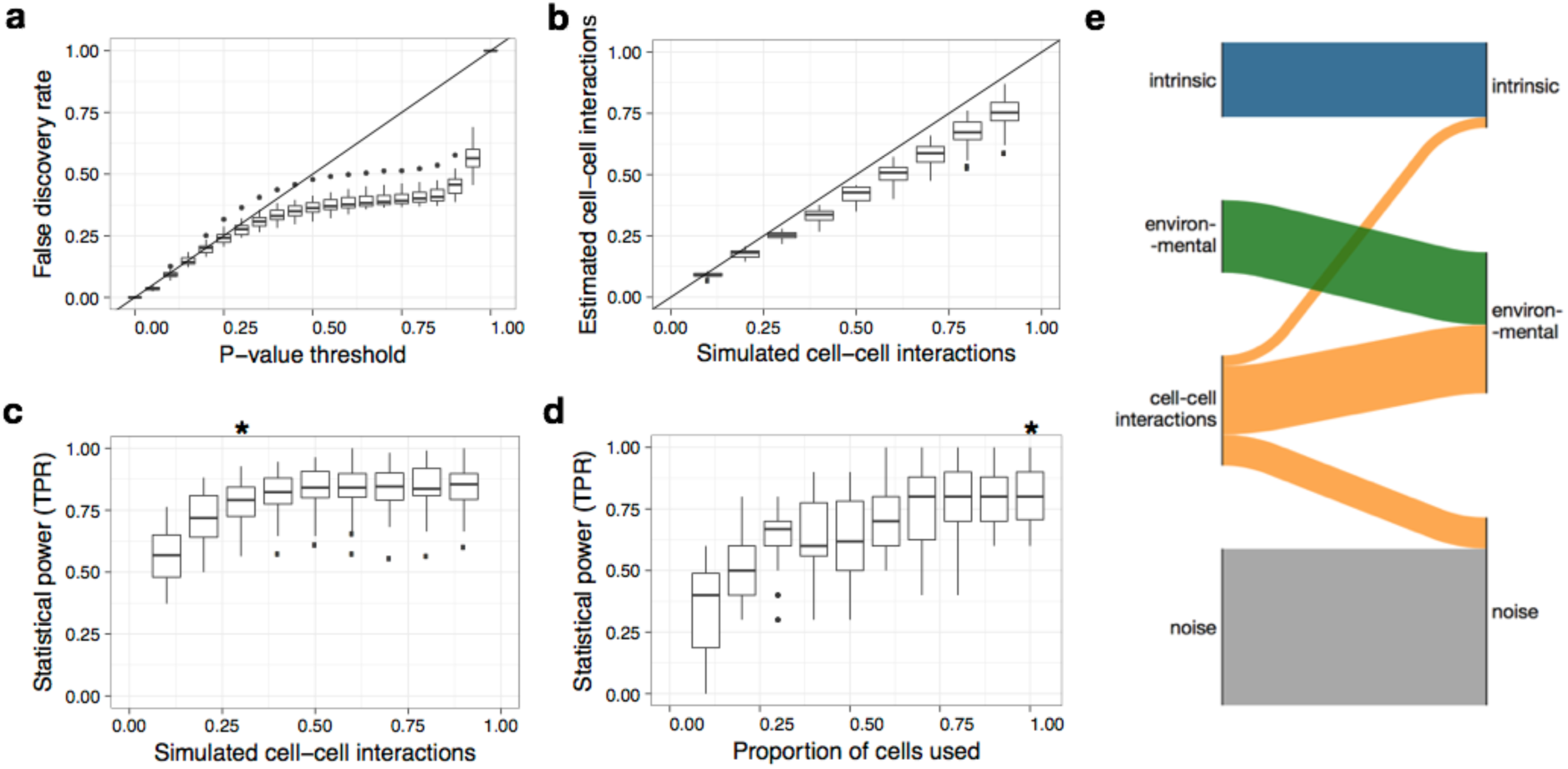
Validation of cell-cell interaction tests and variance components using simulated data. **(a)** Empirical false discovery rate for the cell-cell interaction test using data simulated from the null, without cell-cell interactions. Shown is the empirical false discovery rate (FDR) as a function of the P value threshold. **(b)** Fraction of variance due to cell-cell interactions estimated by SVCA when varying the true fraction of variance explained by cell-cell interactions (**Methods**). **(c,d)** Statistical power of the test for cell-cell interactions (at family-wise error rate <1%), when varying the simulated fraction of variance explained by cell-cell interactions (**c**) and when considering different subsets of cells from the full dataset for model fitting (**d**). The full dataset with all cells was used for model fitting in **c** (indicated using the asterix symbol). In each panel, boxplots display the distribution of results across 26 proteins. Rates in panel **a**, **c**, **d** (True Positive Rate - TPR and False Discovery Rate - FDR) are computed for each protein, aggregated across 110 simulations (11 images times 10 repeat experiments). Similarly, panel **b** depicts average variance estimates across the same set of 110 simulations for each protein. (**e**) Sankey plot displaying how the variance explained by cell-cell interactions is captured by the other terms when omitting interaction effects in the model. Bar height denotes estimated variance fractions (lef: SVCA; right: reduced model). Linking edges indicate the redistribution of variance estimates between the full and the reduced model. Both models were fitted on the same simulated data for 11 images and 26 proteins with cell-cell interactions explaining 30% of the variance of simulated expression levels and results shown are averaged across images and proteins. (**Methods**).

### Application of SVCA to spatial proteomics data of breast cancer tissues

Next, we applied SVCA to an Imaging Mass Cytometry (IMC) dataset from human breast cancer, consisting of 52 breast biopsies from 27 breast cancer patients with variable disease grade and from different cancer subtypes, sampled from different tumour locations (Schapiro et al., 2017). SVCA revealed substantial differences of the overall importance of cell-cell interaction components across proteins, explaining up to 25% of the total expression variance on average (**Fig 3a**, **Supp. Table 1**). Immune cell markers in particular were enriched among the set of proteins with the largest cell-cell interaction effects: CD44, CD20, CD3 and CD68, for which we detected significant cell-cell interaction effects in 35, 34, 34 and 38 out of the 52 images respectively (FDR<1%, Benjamini-Hochberg adjusted, **Fig. 3a**). We used bootstraps to confirm the robustness of the variance estimates (**Fig. S4**), and we observed that a variance component model that accounts for cell-cell interactions yielded more accurate gene expression profiles imputations than models that ignore such effects (**Fig. 3b, Fig. S2)**.

**Figure 3.**
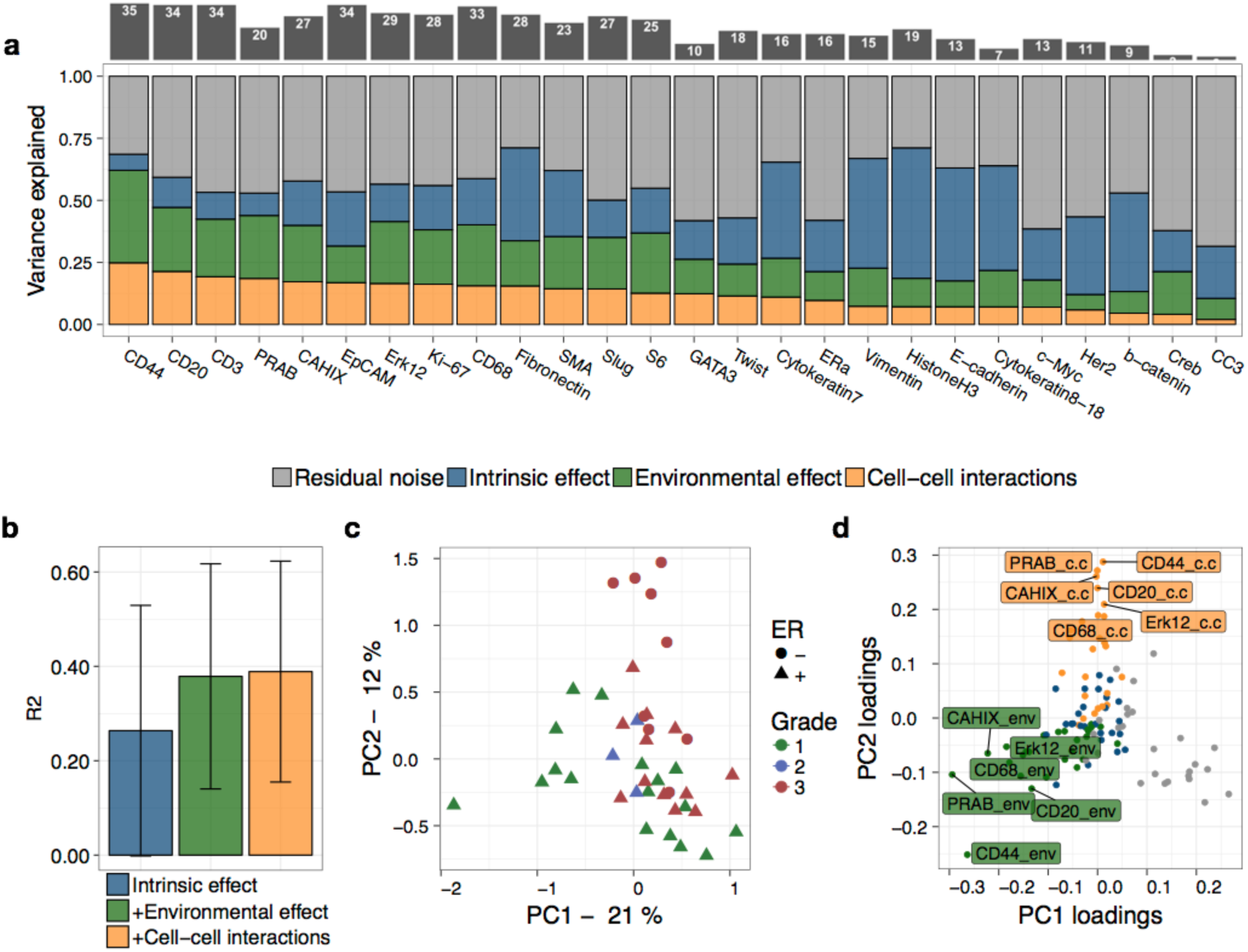
Application of SVCA to 52 breast cancer samples profiled using IMC. (**a**) Bottom panel: SVCA signatures for 26 proteins. Shown are averages of the fraction of variance explained by intrinsic effects, environmental effects and cell-cell interactions, across 52 images. Proteins are ordered by the magnitude of the cell-cell interaction component. Top panel: number of images with significant cell-cell interaction effects for the corresponding proteins, out of 52 images (FDR<1%, **Methods**, **Fig S5**). (**b**) Accuracy of SVCA and models of lower complexity to predict gene expression markers for hold out cells (5 fold cross validation, masking cells). Shown are average coefficients of determination (r2) between predicted and observed gene expression profiles, averaged across proteins and images. Error bars correspond to plus and minus one standard deviation across images and proteins. (**c**) First two principal components for 42 clinically annotated images, calculated based on the spatial variance signature, with individual images coloured by tumour grade. (**d**) Loadings of the principal components as in **c**, depicting the relevance of individual proteins and types of variance components.

We also observed substantial differences of the estimated spatial variance signatures between different images (**Fig. S3**), motivating investigating the relationship between these variations and clinical covariates, including tumour grade, ER status, PR status, HER2 status. A projection of the full SVCA output using principal component analysis (PCA) identified tumour grade as the most important explanatory variable for differences in spatial variance components (**Fig 3c**), followed by ER status and PR status (**Fig. S6**). Inspection of the PCA loadings (**Fig. 3d**) identified the cell-cell interaction component and the environmental component for a subset of proteins (including PR and CD44) as the most informative SVCA features affected by tumour grade. This was also evident when comparing environmental and cell-cell interaction effects across tumours of different grades (**Fig. S7**), showing a relative increase in the cell-cell interaction component in higher grade tumours. This effect could partially be linked to the underlying disorganized and irregular cellular architecture characterizing these high grade tumors, which is associated with larger cells, increased proliferation and thus potentially higher cell density in comparison to healthy breast tissue (Elston & Ellis, 1991).

### RNA datasets based on imaging technologies

SVCA can be used for the analysis of data from a broad range of spatially resolved technology, including optical imaging-based assays. To explore this, we considered a mouse hippocampus datasets, profiling 249 RNA expression levels in 21 distinct brain regions of a single animal, using seqFISH (Shah et al., 2017). Spatial Variance signatures for all genes are given on **Supp. Table 2** and on **Fig. 4a** for the top 20 genes for cell-cell interactions. Leveraging the higher dimensionality of these data, we sought to identify individual genes and pathways that participate in cell-cell interactions.

**Figure 4:**
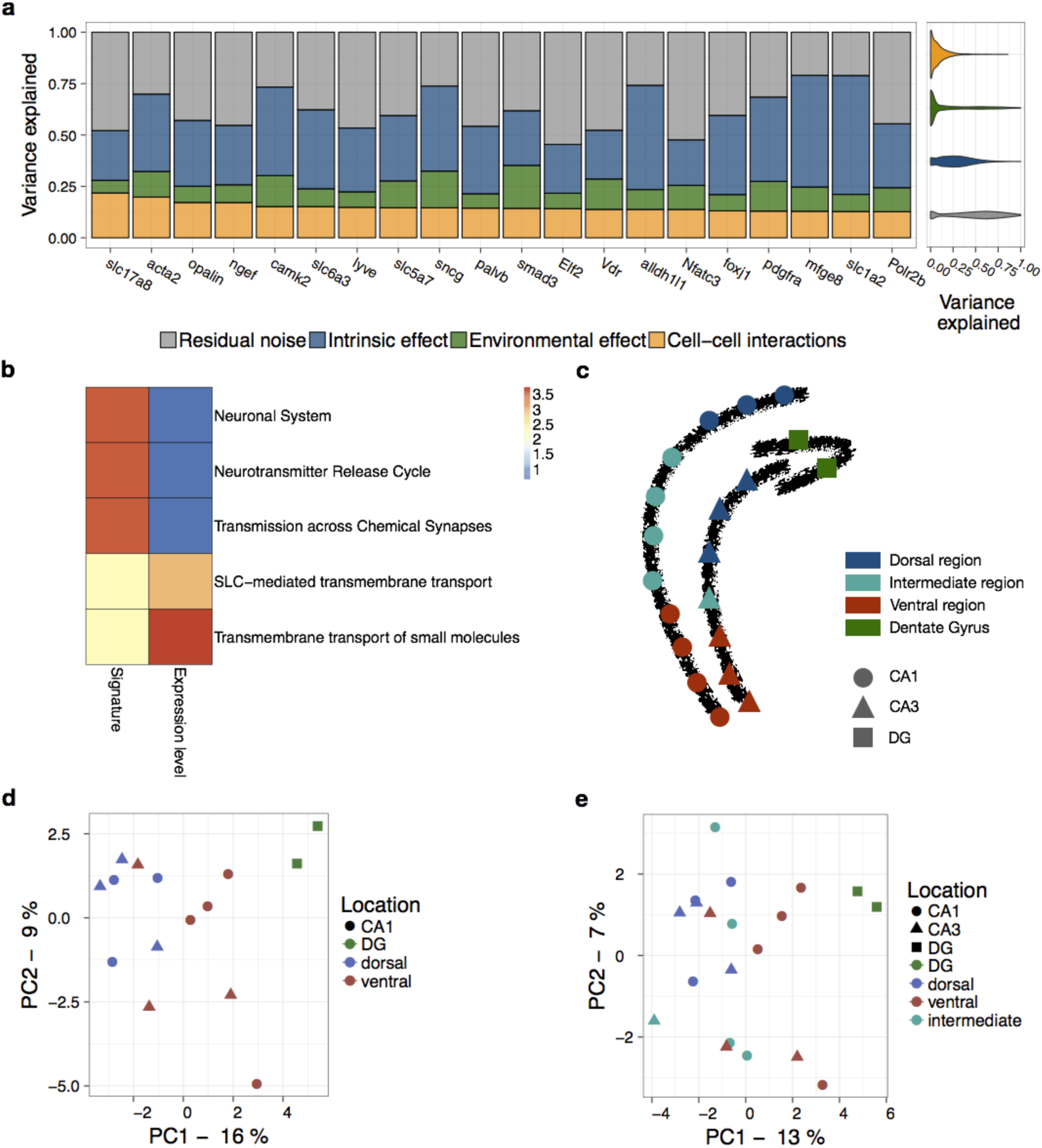
Application of SVCA to 20 images profiled using seqFISH. **(a)** Left: SVCA signatures for the top 20 proteins with highest cell-cell interaction effects. Shown are averages of the fraction of variance explained by intrinsic effects, environmental effects and cell-cell interactions, across 20 images. Proteins are ordered by the magnitude of the cell-cell interaction component. Right: Variance estimates distribution across images and proteins for all genes (violin plots). **(b)** Top 5 enriched pathways out of 80 Reactome pathways considered using a rank-based enrichment test (based on a consensus ranking across images, **Methods**), either considering the fraction of variance explained by cell-cell interactions (left), or the expression level of individual genes (right). Colours denote statistical significance (negative log Benjamini-Hochberg adjusted P values). **(c)** Spatial organisation of the mouse hippocampus with dots corresponding to individual images. Colours and shapes denote regions using the classification as in (Shah et al., 2017). **(d)** First two principal components of the spatial variance signatures for individual images from the DG, the dorsal region and the ventral region. Colour and shape represents the location of the biopsy in the hippocampus. **(e)** First two principal components of the spatial variance signatures for all 20 images.

Similarly to the IMC dataset, SVCA signatures were robust and could accurately predict gene expression levels out of sample (**Fig S13**, **S14**, **S15**). We considered the average cell-cell interaction components across images and tested for enriched Reactome pathways (Croft et al., 2014), (**Methods**). This identified five pathways with significant enrichments (FDR<5%, considering 80 pathways with five or more genes contained in our dataset, **Methods**, **Fig. 4b**, **Supp. Table 3**), with *transmission across chemical synapses, neurotransmitter release cycle* and *neuronal system* being among the most significant ones. For comparison, we also considered gene set enrichments based on the average gene expression levels across images, which identified distinct enriched pathways (*transmembrane transport* and *SLC-mediated transmembrane transport*). These results demonstrate that spatial and abundance-based signatures yield complementary information for studying biological activity in tissues.

Similarly to results obtained on the IMC datasets, we observed differences in the spatial variance signatures across images, which were sampled from functionally distinct regions of the hippocampus (Shah et al., 2017). Principal components of the spatial variance signature for the dorsal region clustered together, irrespective of their CA1/CA3 location. Similarly, images from the Dentate Gyrus (DG) also clustered together, and there was some proximity between signatures from the ventral region, although with more variation between them (**Fig. 4d,e**). This is consistent with Shah et al’s observation that the ventral and dorsal regions of the CA1 and the CA3 mirrored each other with respect to their cellular compositions, and that ventral regions are more heterogeneous in their cellular composition. Spatial variance signatures for intermediate regions, however, did not show much resemblance (**Fig. 4d,f**).

Finally, we applied SVCA to a breast cancer cell culture dataset profiled using the mer-FISH technique (130 RNA expression levels, **Supp. Table 4**) (Moffitt et al., 2016). The large spatial domain and the total number of cells imaged (more than 50,000) from a biological homogenous cell culture system enabled defining different field of views as biological replicates (N=20, 2,600 cells on average). Consistent with this, the variance components of SVCA were more consistent across images (**Fig. S8**, **Fig. S9**, **Fig. S10**, **Fig. S12**), with average coefficients of variation for cell-cell interactions ranging between 20 and 40%, compared to typically 75-150% (**Fig. S11**) for the results obtained from the IMC and seq-fish data (**Fig. S3**, **Fig. S16**). These results further support the general applicability of the model to data from different technologies and the robustness of SVCA variance signatures.

## Conclusion

We presented Spatial Variance Component Analysis (SVCA), a regression-based framework for the analysis of spatially resolved molecular expression data. Our model computes a spatial variance signature for individual mRNA or protein levels, decomposing their sources of variation into spatial and non-spatial components. Most prominently, SVCA provides a quantitative assessment of the effect of cell-cell interactions on the expression profile of individual molecules. The model avoids the explicit definition of cell types and neighbourhoods, instead using continuous measure of cell state and euclidean distances between cells.

We have applied SVCA to multiple datasets generated using alternative technologies, probing either RNA transcripts or proteins, demonstrating the broad applicability of the model. Across these applications, we observed that cell-cell interactions can substantially contribute to gene expression variation, which is consistent with previous reports (Battich et al., 2015; Goltsev et al., 2018; Kamińska et al., 2015; Nasra Naeim Ayuob and Soad Shaker Ali, 2012) and supports the concept that studying single cell expression in the native context is important for understanding the sources of these variations. We also showed that the variability of spatial variance signatures across samples of the same biological system could be linked to different clinical contexts or to the internal structure of a given organ, which underlines the biological relevance of these signatures. Finally, using higher dimensionality data obtained from optical technologies, we observed that genes with largest cell-cell interaction components were enriched in specific pathways, suggesting that spatial variance signatures could help us understanding biological activity in tissue. These results suggest that analysing single cell data in their native context, with SVCA or related methods, will help us understand how the spatial organisation of tissues impact single cell biology.

Although we have tested the calibration and robustness of SVCA, the model is not free of limitations. At present, the model does not account for technology-specific noise and instead assumes Gaussian distributed residuals, thus requiring suitable processing of the raw data such that these assumptions are sufficiently met (See **Methods**). Further development could consider a generalized random effects model, for example to couple the random effect component with a negative-binomial likelihood. A second limitation of SVCA is that the model is univariate, which means that individual genes or proteins are modelled independently from each other. Multivariate extensions could account for relationships between genes involved in the same pathways, either in an unsupervised manner or using prior knowledge (Buettner, Pratanwanich, McCarthy, Marioni, & Stegle, 2017). Such approaches could give a more comprehensive understanding of how biological processes are affected by tissue structure.

There is a growing appreciation of the role of spatial distribution of proteins, transcripts and other molecules in determining tissues functioning and its deregulation in disease, with potential value as predictors of clinical outcomes (Bodenmiller, 2016). This is largely driven by vigorous development of novel technologies that enable us to capture such data (Aichler & Walch, 2015; Bodenmiller, 2016; Goltsev et al., 2018; Lin et al., 2017; Schulz et al., 2018). We believe that the SVCA framework and extensions thereof will be of broad use to analyze this burgeoning spatially-resolved molecular data to advance our understanding of the pathophysiology of multiple diseases.

## Methods & software

A full derivation of SVCA and details on the analysis are provided as Supplementary Methods. An open source implementation of SVCA is available at https://github.com/damienArnol/svca.git, which builds on the limix package (Lippert, Casale, Rakitsch, & Stegle, 2014).

## Author contributions

Author contributions: DA, OS developed the statistical method. DA implemented the model and analysed all the data. DS, BB contributed to the interpretation of the data and the design of the method. DA, DS, OS, JSR wrote the manuscript with input from all authors. JSR and OS conceived the project and supervised the work.

## Acknowledgments

DA, JSR, OS acknowledge EMBL core funding. DS was supported by the Forschungskredit of the University of Zurich, grant FK-74419-01-01, and the BioEntrepreneur-Fellowship of the University of Zurich, reference no. BIOEF-17-001. B.B.’s research is funded by an SNSF R’Equip grant, an SNSF Assistant Professorship grant, the SystemsX Transfer Project “Friends and Foes”, the SystemsX MetastasiX and PhosphoNetX grant, NIH grant (UC4 DK108132), and the European Research Council (ERC) under the European Union’s Seventh Framework Program (FP/2007-2013)/ERC grant agreement no. 336921.

We thank R. Argelaguet, V. Svensson, R. Vento, H. Jackson, A. Baud, N. Cai, F.P. Casale and D. Horta for discussions on data processing, model design and implementation and results visualisation.

## Supplementary Figures

**Figure S1.**
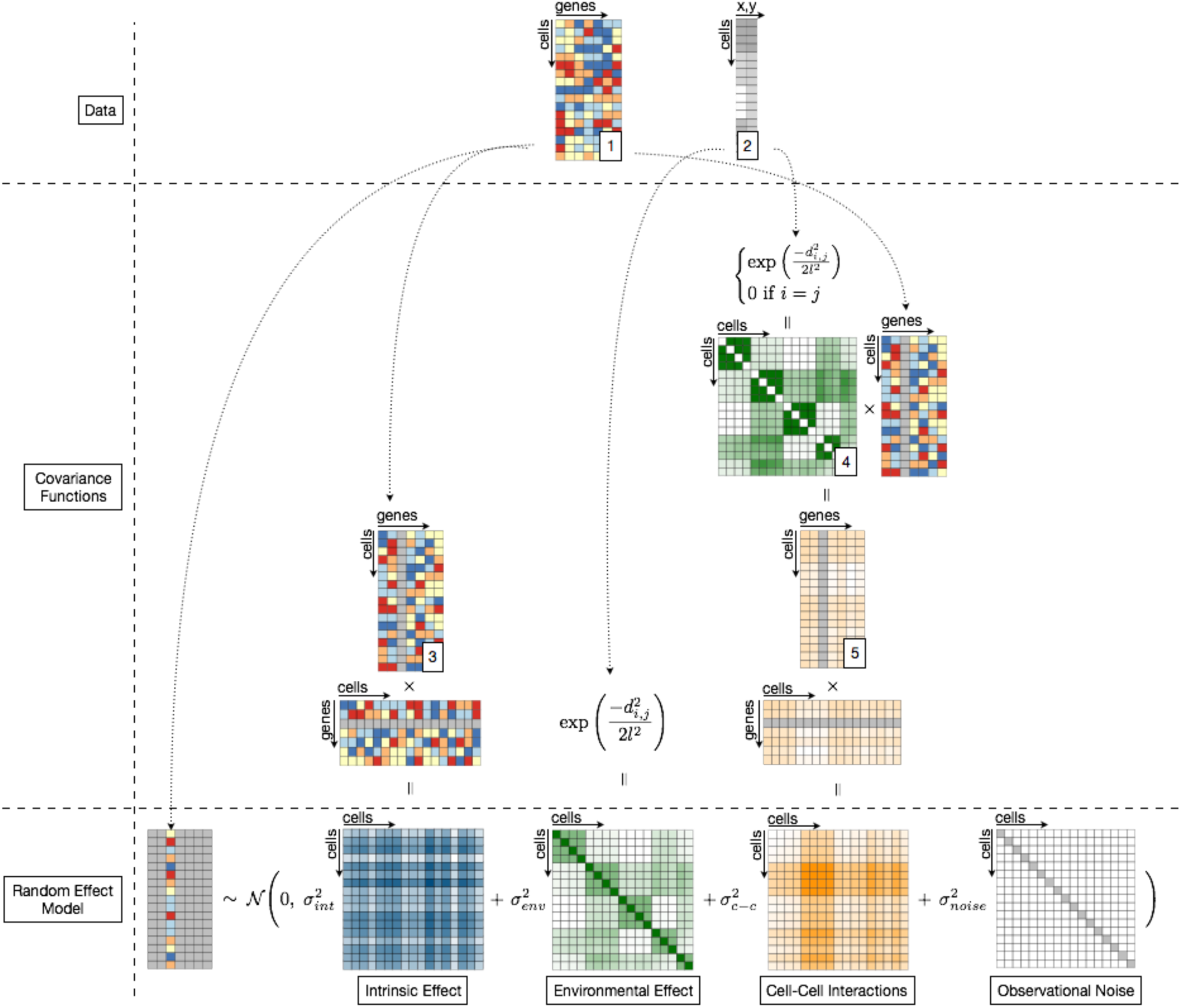
SVCA model definition. SVCA takes as an input single cell expression data as a cell times gene/protein matrix (1) and the spatial location of the cells as a cell times (x,y) coordinates matrix (2). Individual genes are modelled as multivariate normally distributed, with additive covariance components that account for different effects modelled by SVCA (intrinsic, environmental and cell-cell interactions, Fig.1). The covariance for intrinsic effects is computed as the empirical covariance of the expression profiles between cells, where the modelled gene has been removed from the expression matrix (3). Environmental effects are modelled using a Squared Exponential covariance defined on the relative distance between cells. Cell-cell interactions are modelled using a cellular neighbourhood matrix (5) which aggregates, for each cell, the molecular composition of the neighbouring cells. This is achieved by weighting the molecular profiles of all neighbouring cells with a squared exponential function of the distance. This can be written as a product between a squared exponential covariance matrix whose diagonal elements were removed (4) and the expression matrix (3). The final cell-cell interaction covariance is computed as the empirical covariance of the cellular environment matrix between cells. SVCA’s training then consists in learning the scale of every covariance term.

**Figure S2.**
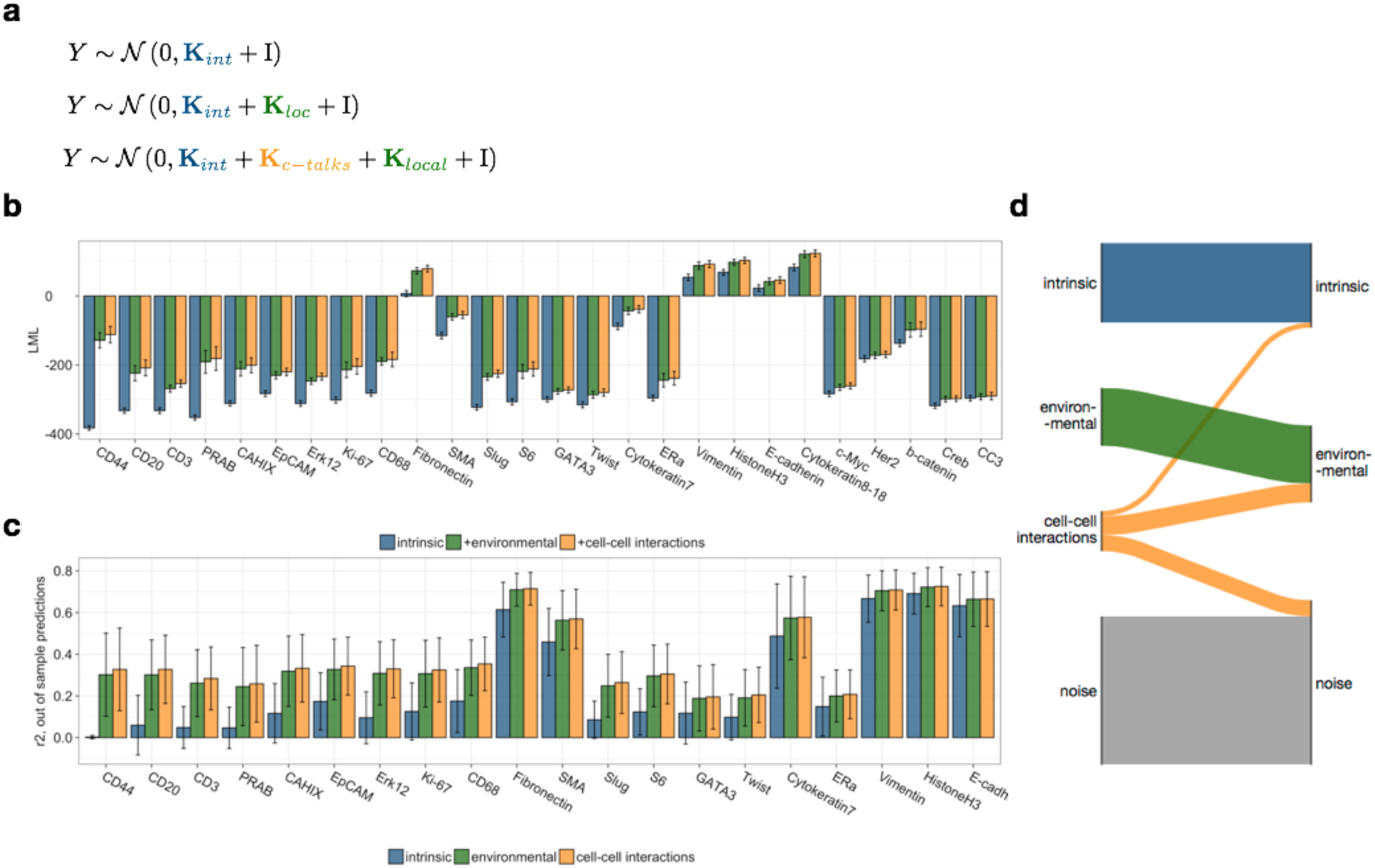
Assessment of the predictive performance of SVCA and reduced models. **(a)** Considered here are three models in increasing order of complexity. The first model accounts only for the intrinsic component, the second model also includes the environmental effect. The third model includes all three components of SVCA **(b)** Log marginal likelihood of the three models of increasing complexity **(c)** Out of sample prediction performance (r2) for the three models of increasing complexity (5-fold cross validation, **Methods**). The results show that the increase in complexity also improves the generalisation performance to a validation set. Proteins ordered by the size of the cell-cell interaction component. **(d)** Variance redistribution from the SVCA model to a reduced model including intrinsic and environmental effects only, averaged across all proteins (see Fig. 2).

**Figure S3.**
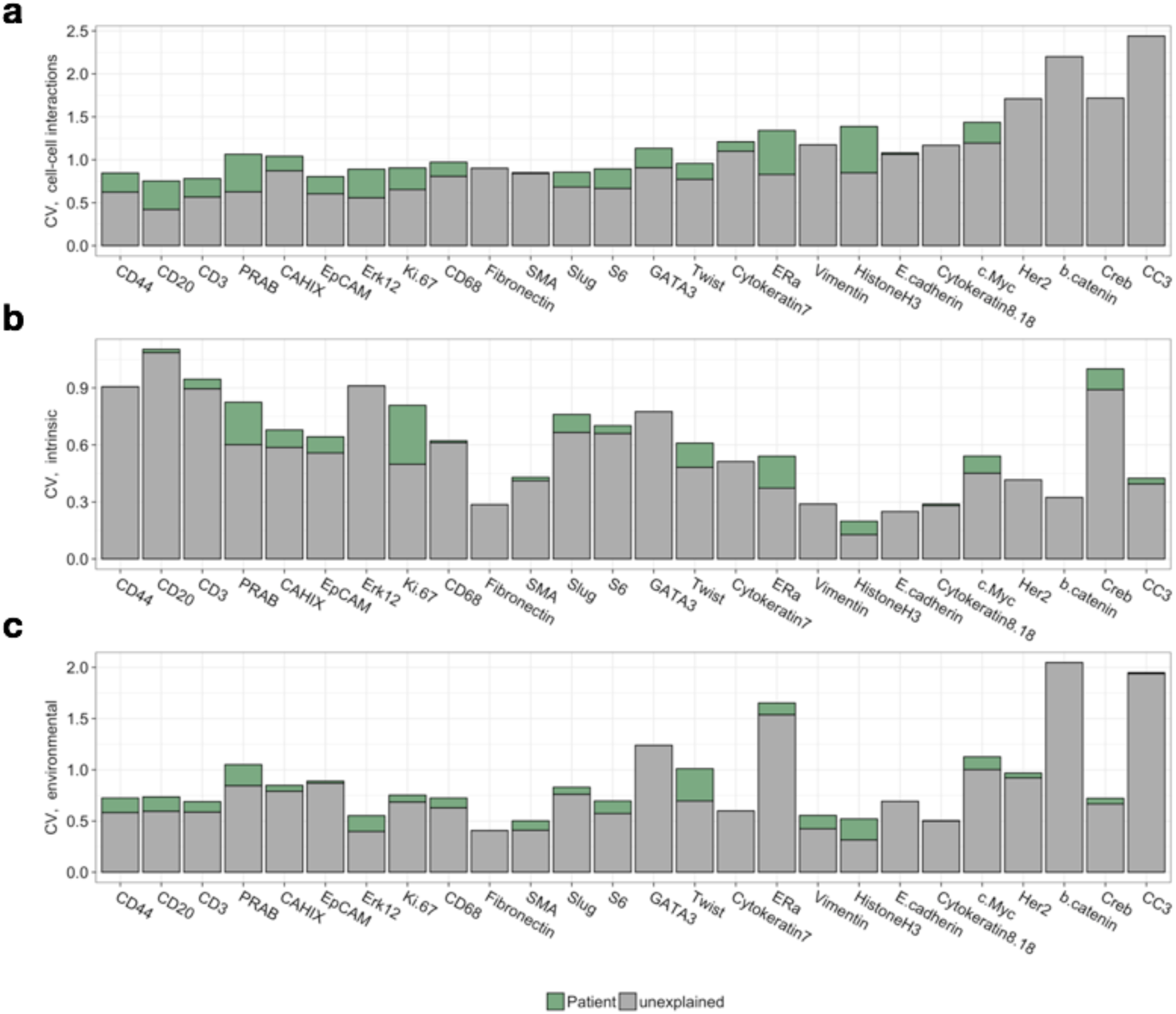
Coefficient of variation of SVCA variance components across samples. Coefficient of variation of the different variance components. Shown in green is the proportion that is attributable to patient, estimated using a random effect model (see **Methods**) **(a)** Cell-cell interaction component **(b)** Intrinsic component **(c)** Environmental component.

**Figure S4.**
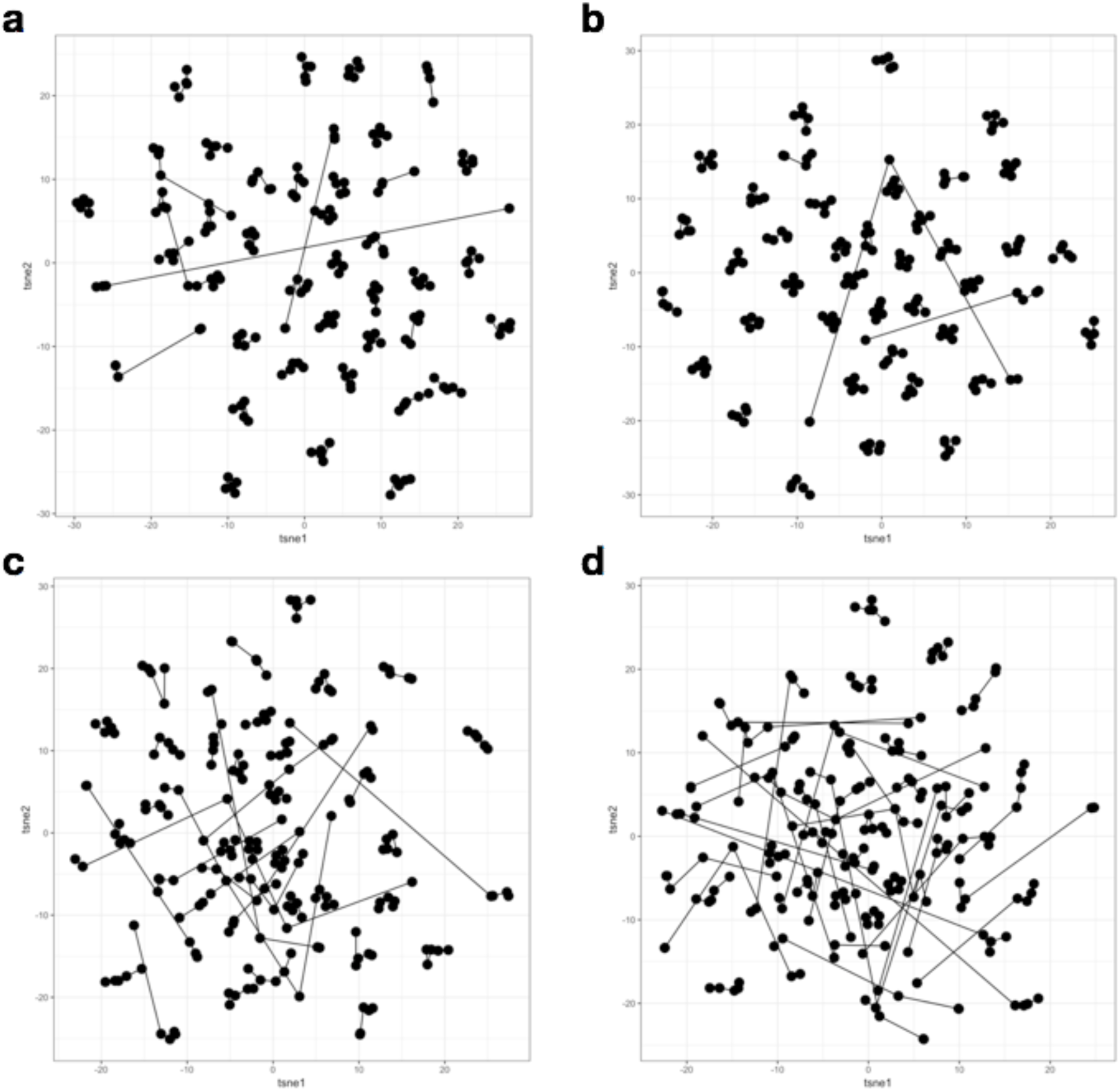
Spatial variance signature robustness using a bootstrapping strategy (see Methods). t-SNE representation of the spatial variance signature components including **(a)** the entire SVCA signature, **(b)** only the intrinsic component, **(c)** only the cell-cell interaction component **(d)** only the environmental component. Results from multiple bootstraps for the same image (linked with a line) cluster together, confirming the robustness of the spatial variance signatures

**Figure S5.**
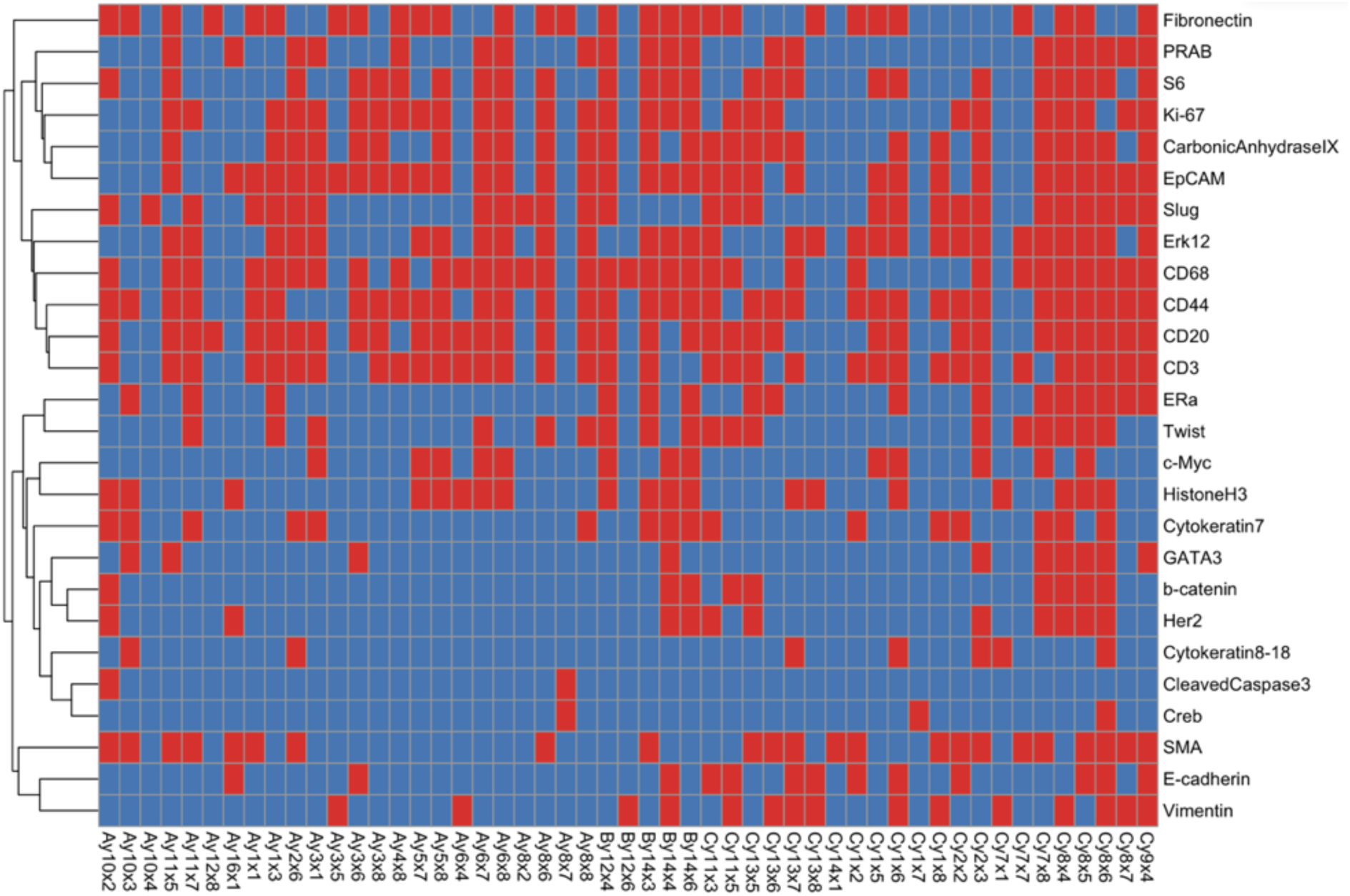
Significance of the cell-cell interaction components per protein and image. In red, proteins for which the cell-cell interaction component is significant (FDR < 1%, Benjamini-Hochberg adjusted, **Methods**)

**Fig S6.**
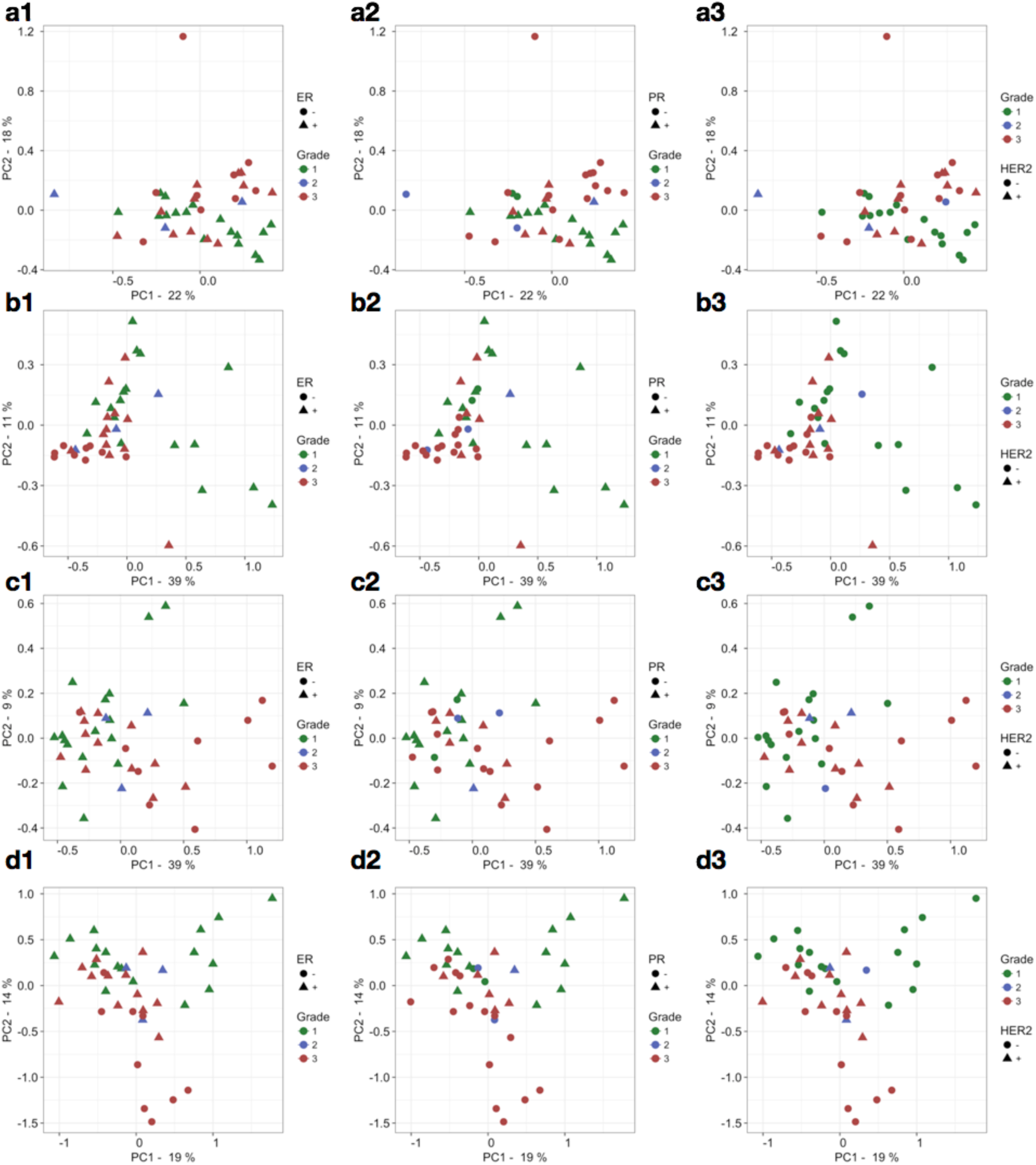
Variance signatures PCA and the relationship with clinical covariates. **(a)** First two components for the intrinsic variance term for all proteins. The colour corresponds to the grade and the shapes correspond to **(a1)** the ER status **(a2)** the PR status and **(a3)** the Her2 status of the samples respectively. **(b)** analogous PCs of the environmental variance component, **(c)** analogous PCs of the cell-cell interaction component and **(d)** Analogous PCs of the entire Spatial Variance signature.

**Fig S7.**
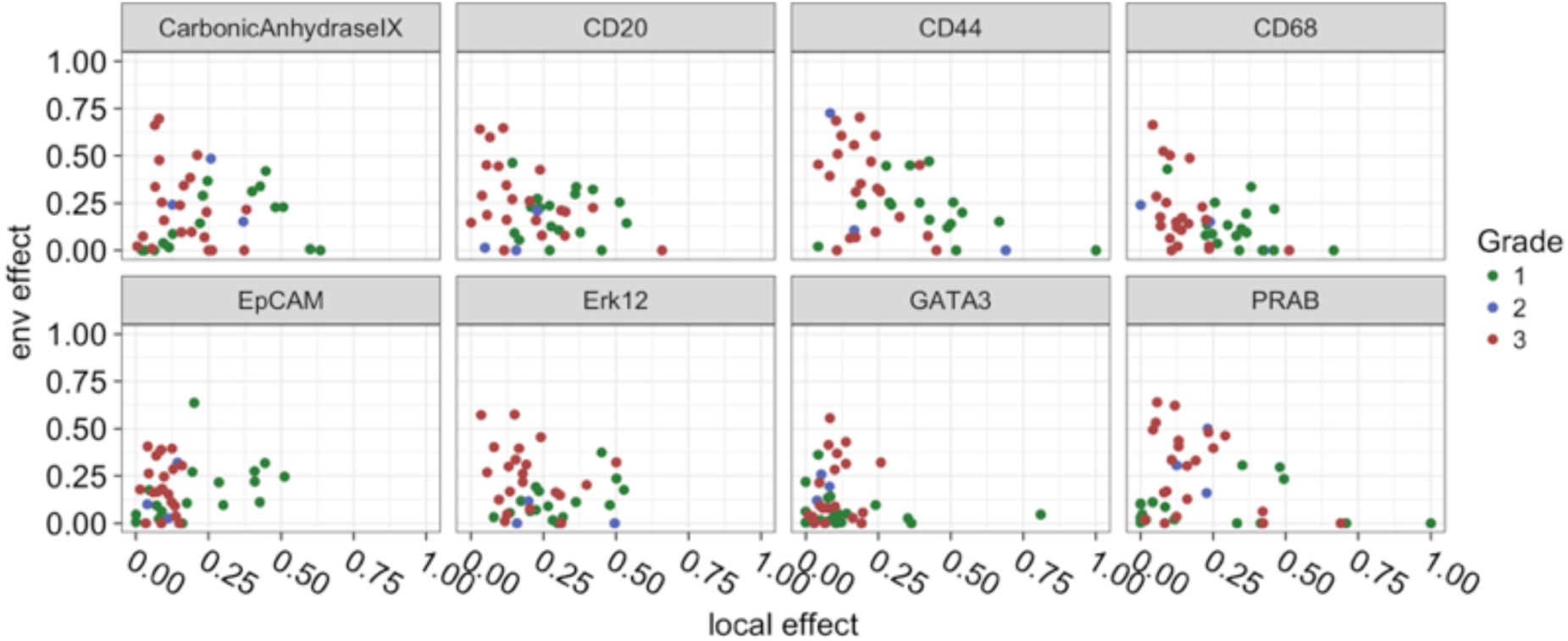
Ratio between cell-cell interaction effects and environmental effect. Proteins are the relevant proteins from the PCA loadings inspection. x axis: environmental effect, y axis: cell-cell interaction effect. Each point corresponds to an image and the colour corresponds to the cancer grade

**Fig S8.**
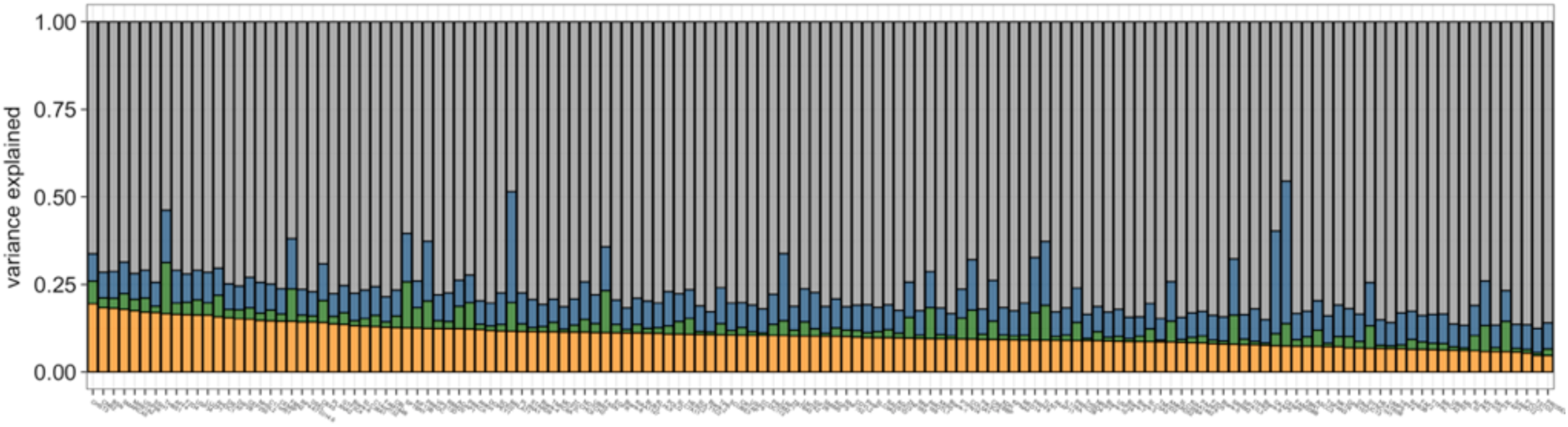
Application to mer-FISH data. Spatial variance signature for 130 genes ordered by the estimated fraction of variance attributable to cell-cell interactions.

**Fig S9.**
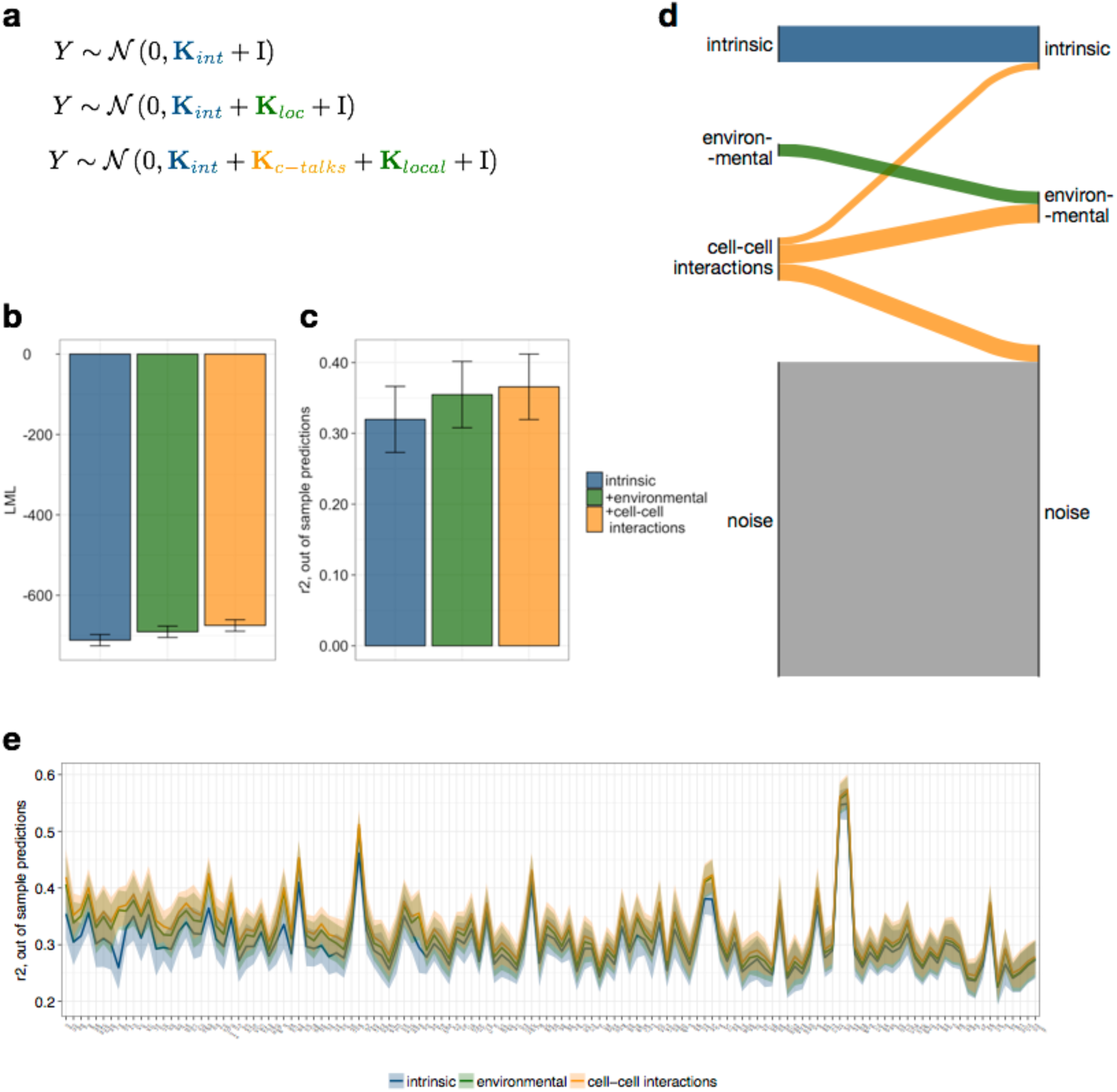
Model comparison mer-FISH data. **(a)** Models compared in order of increasing complexity **(b)** Log marginal likelihood of the three models in increasing order of complexity. Results averaged across all genes. **(c)** Out of sample prediction: coefficient of determination averaged across all proteins for the models in increasing order of complexity. **(d)** Variance redistribution from the SVCA model to a reduced model including intrinsic and environmental effects only, averaged across all proteins (see **Fig. 2**). **(e)** Out of sample prediction for the three models, all proteins. Shaded areas correspond to plus and minus one standard deviation.

**Fig S10.**
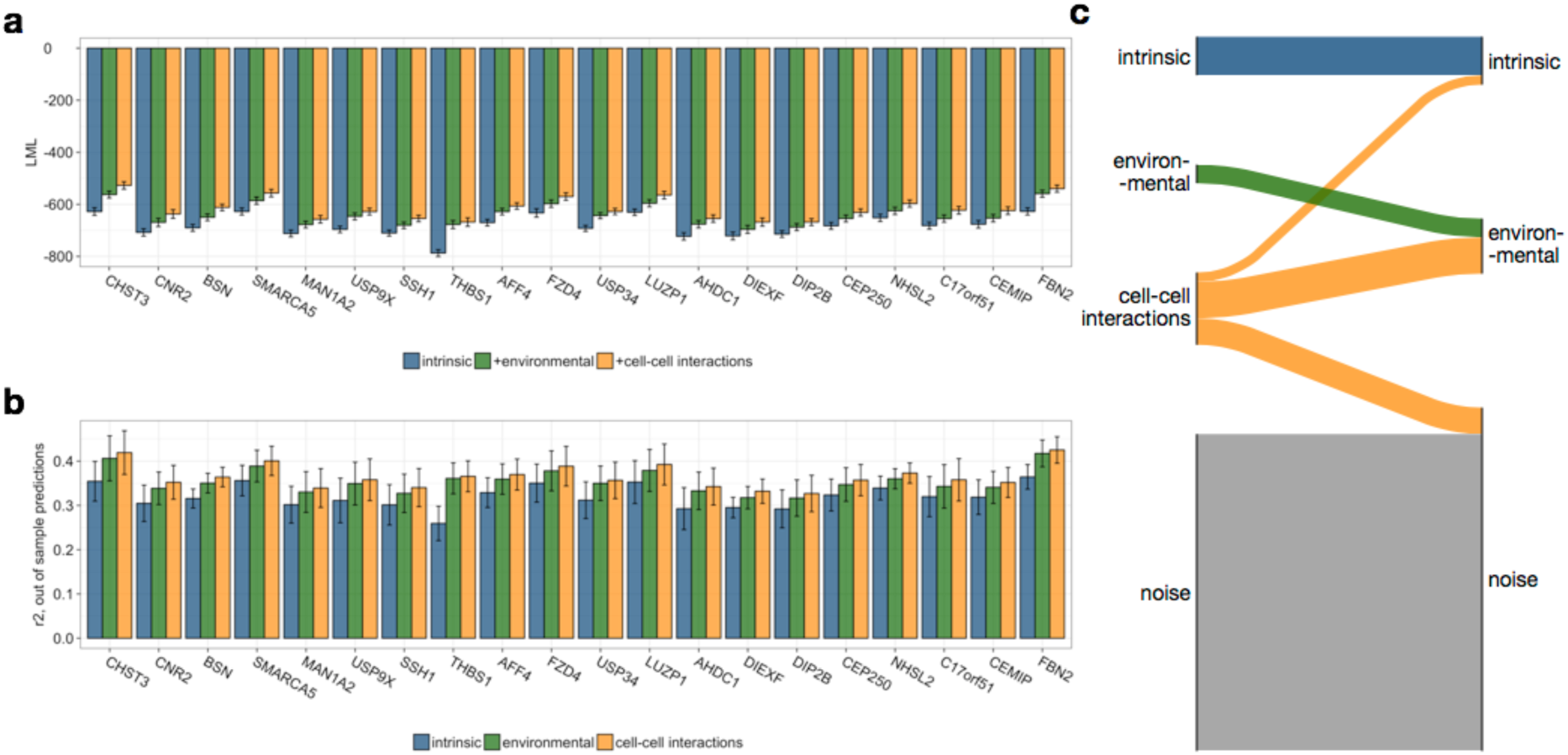
Model comparison mer-FISH data, top 20 cross talk genes. **(a)** Log Marginal Likelihood **(b)** Out of sample prediction r2 **(c)** Variance redistribution from the SVCA model to the reduced model omitting cell-cell interactions (see **Fig. 2**).

**Fig S11.**
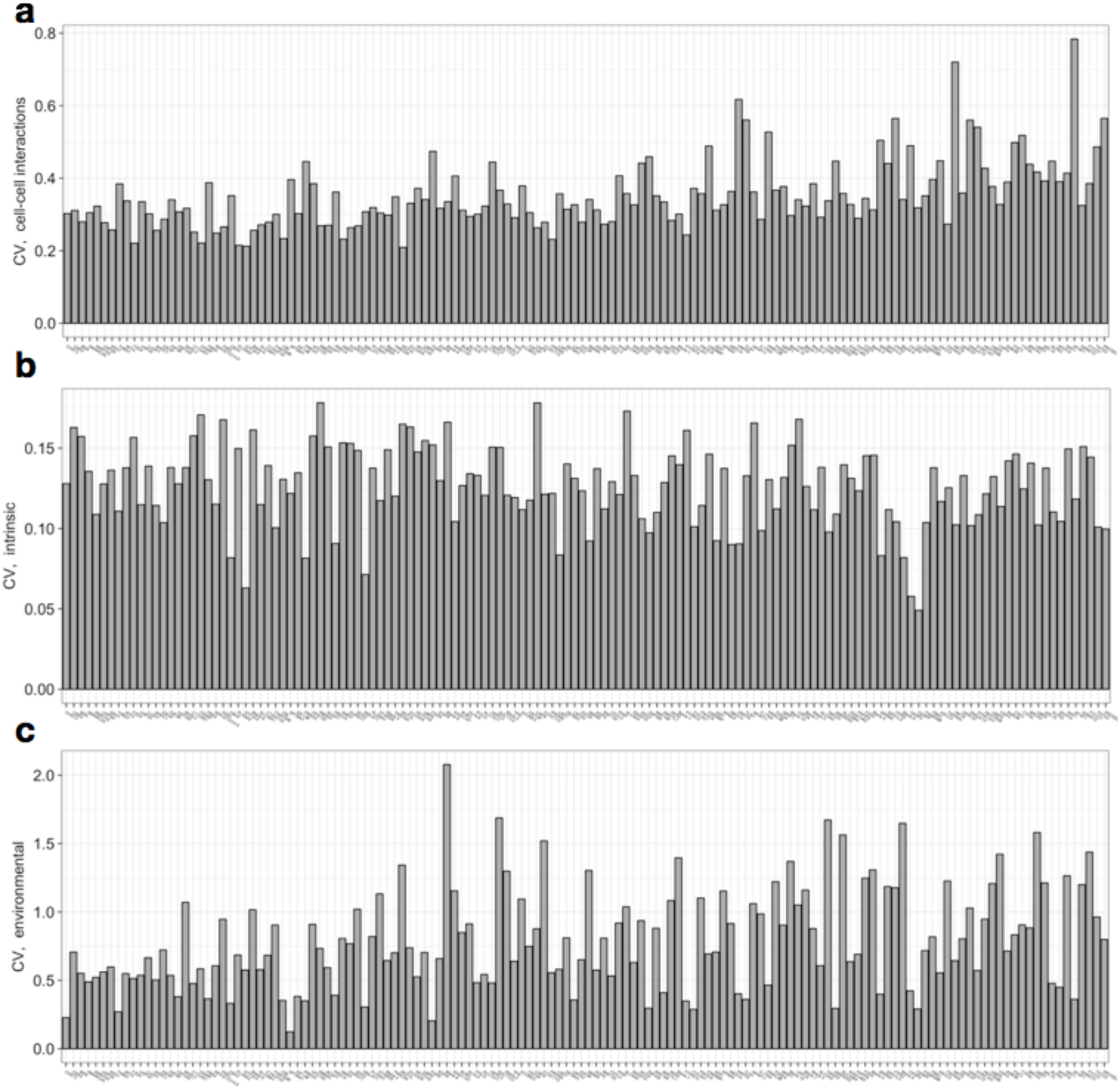
Coefficient of variation of variance components (mer-FISH data). **(a)** Cell-cell interaction term **(b)** Intrinsic term **(c)** Environmental term.

**Fig S12.**
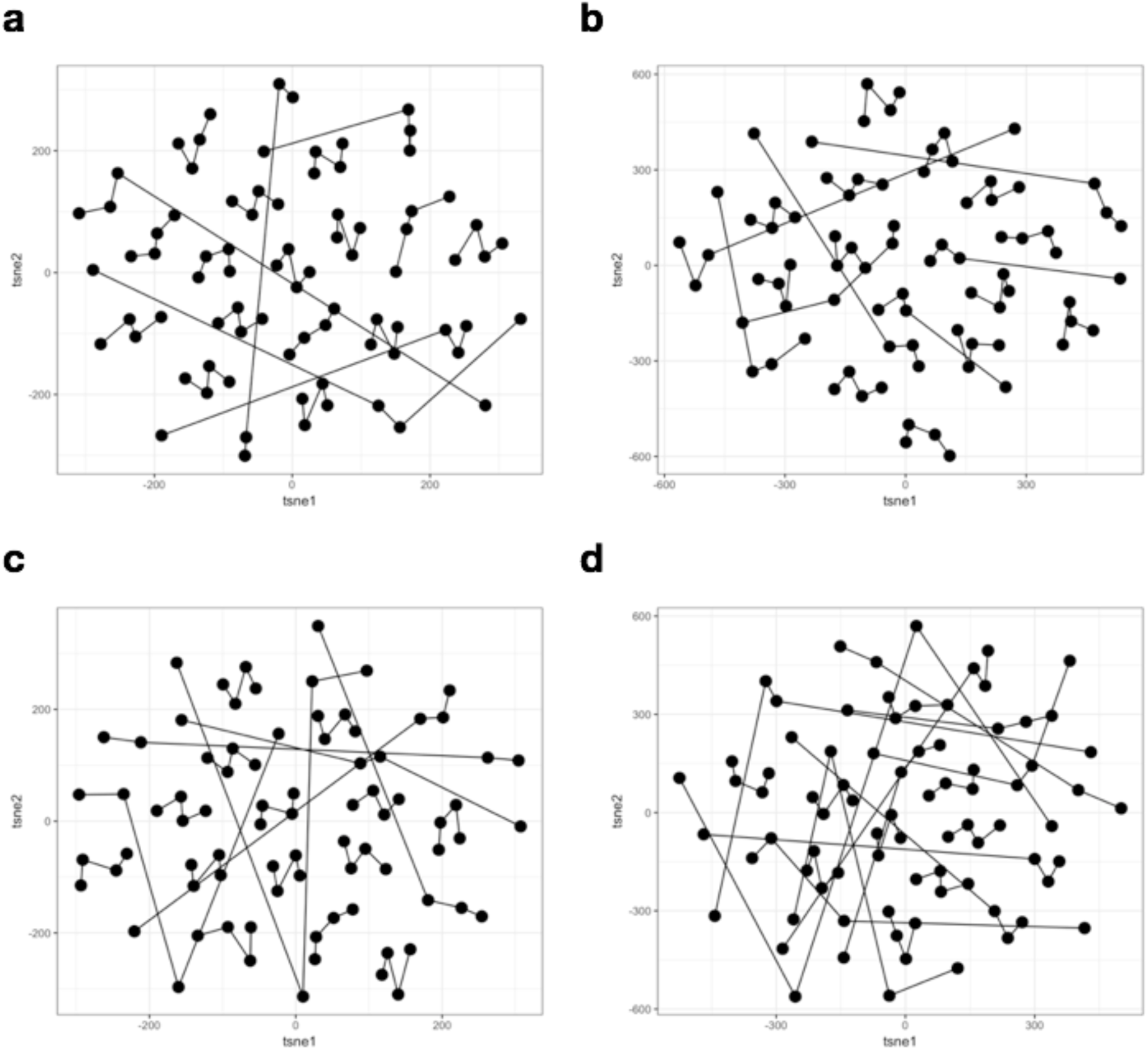
Signature robustness visualised with t-SNE (mer-FISH data). t-SNE representation of the spatial variance signature components including **(a)** the entire SVCA signature, **(b)** only the intrinsic component, **(c)** only the cell-cell interaction component **(d)** only the environmental component. Results from multiple bootstraps for the same image (linked with a line) cluster together, confirming the robustness of the spatial variance signatures.

**Fig S13.**
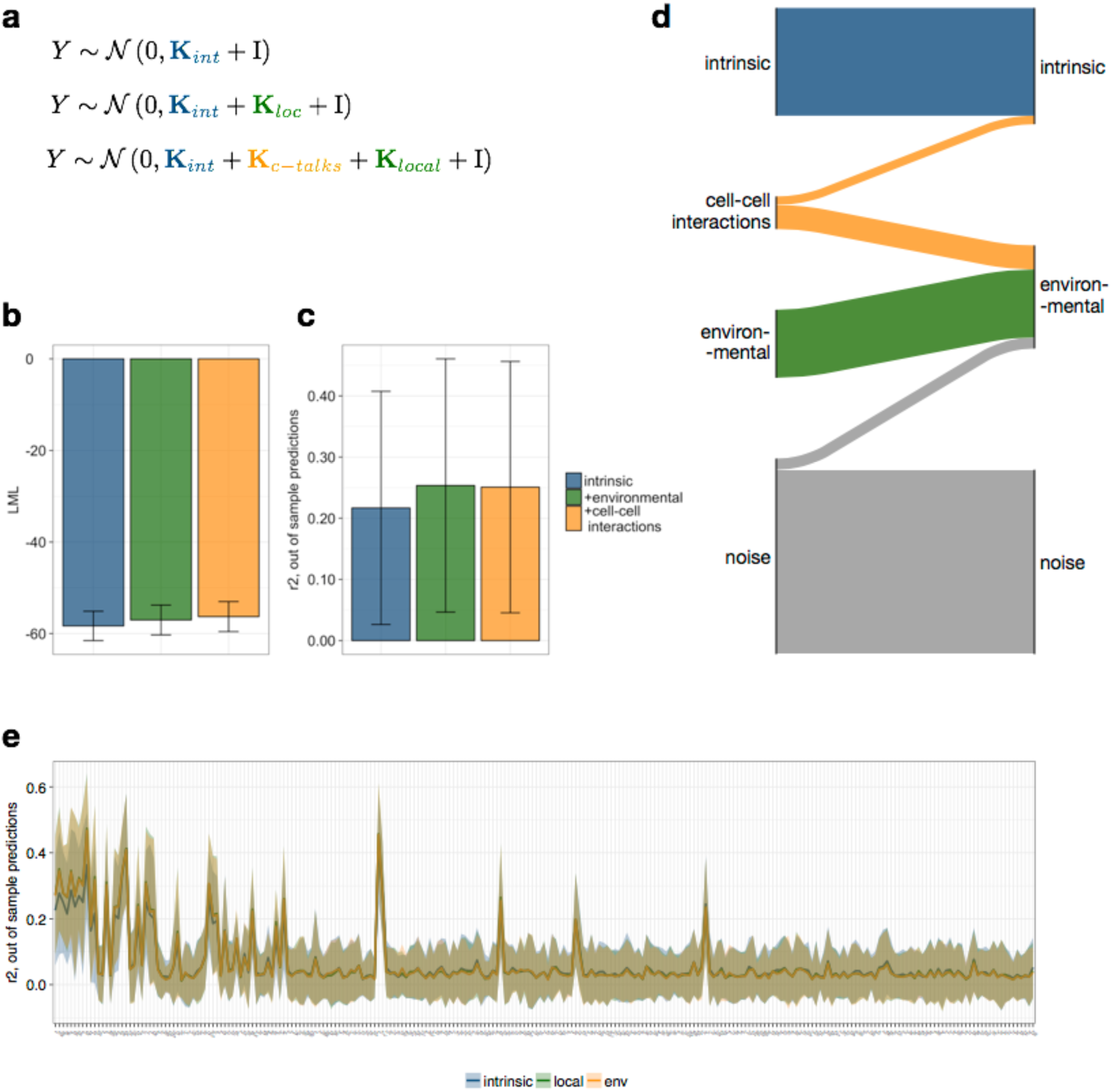
Model comparison seq-FISH data. **(a)** Models compared in order of increasing complexity **(b)** Log marginal likelihood of the three models in increasing order of complexity. Results averaged across all genes. **(c)** Out of sample prediction: coefficient of determination averaged across all proteins for the models in increasing order of complexity. **(d)** Variance redistribution from the SVCA model to a reduced model including intrinsic and environmental effects only, averaged across all proteins (see **Fig. 2**). **(e)** Out of sample prediction for the three models, all proteins. Shaded areas correspond to plus and minus one standard deviation.

**Fig S14.**
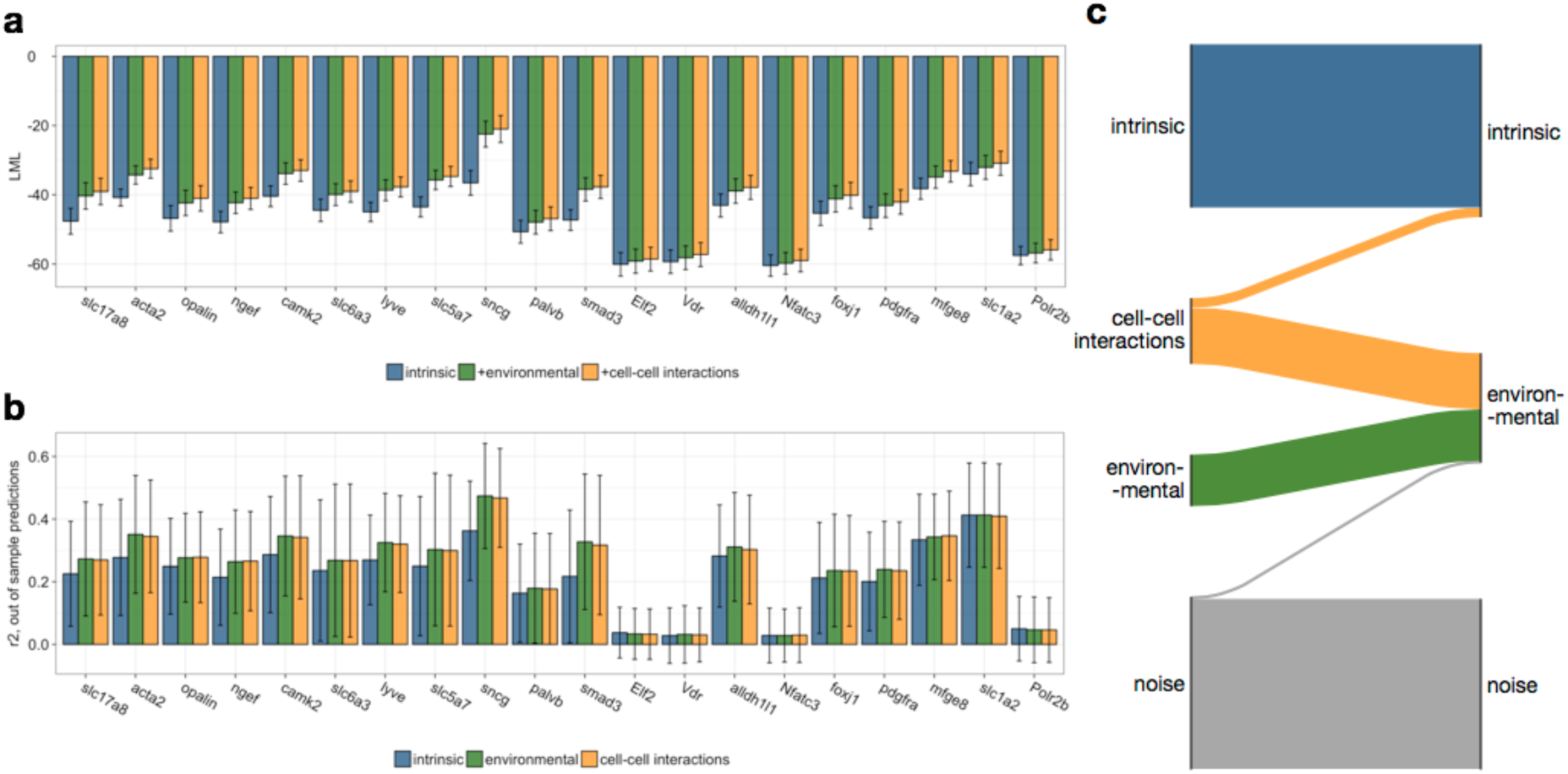
Model comparison seq-FISH data, top 20 cross talk genes. **(a)** Log Marginal Likelihood **(b)** Out of sample prediction r2 **(c)** Variance redistribution from the SVCA model to the reduced model omitting cell-cell interactions (see **Fig. 2**).

**Fig S15.**
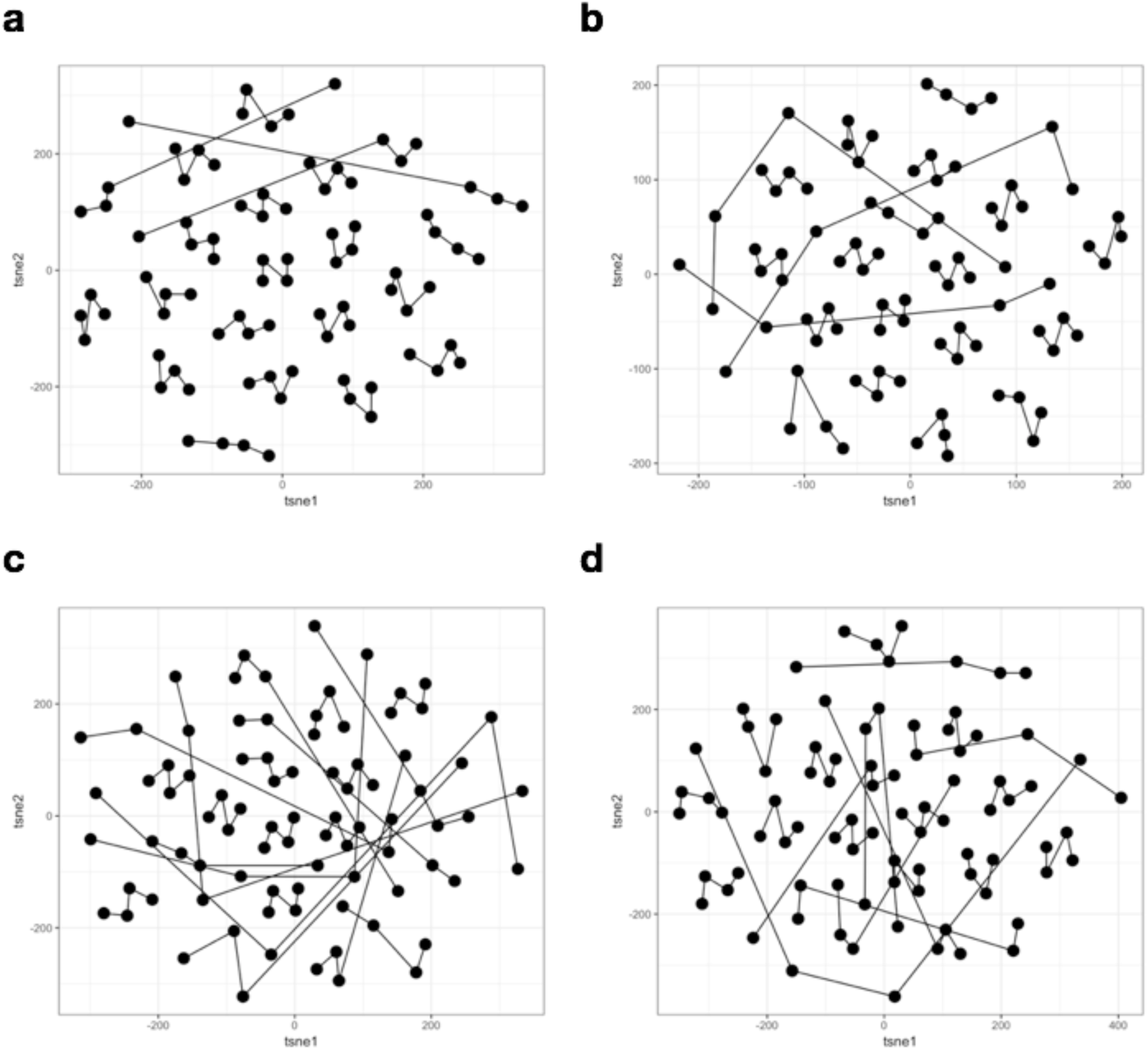
Signature robustness visualised with t-SNE seq-FISH data) t-SNE representation of the spatial variance signature components including **(a)** the entire SVCA signature, **(b)** only the intrinsic component, **(c)** only the cell-cell interaction component **(d)** only the environmental component. Results from multiple bootstraps for the same image (linked with a line) cluster together, confirming the robustness of the spatial variance signatures.

**Fig S16.**
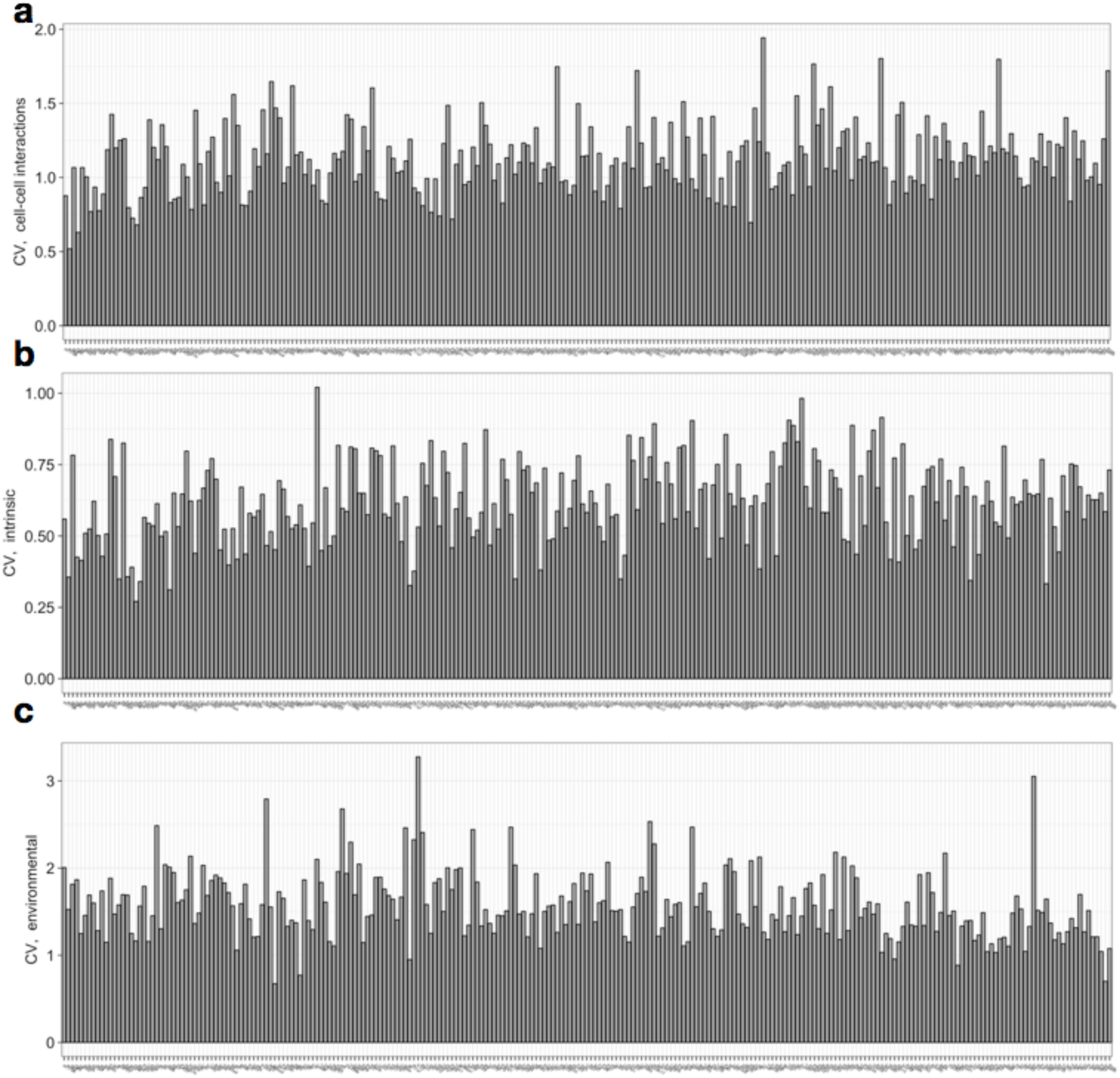
Coefficient of variation of variance components (seq-FISH data) **(a)** Cell-cell interaction term **(b)** Intrinsic term **(c)** Environmental term

## 1 SVCA Model

### 1.1 Model overview

The expression level of a single gene across cells is modelled using a random effect model implemented in a Gaussian Process (GP) framework [16, 13] with an additive covariance. The covariance is composed of 4 terms, modelling our assumption that the variance across cells of the gene expression level is due to three additive effects: an intrinsic effect due to the cell state, a cell-cell interaction effect, due to the state of the neighbouring cells, and an environmental effect due to unobserved factors in the cell micro environment, such as local access to oxygen, nutriments etc.

In the following, we will make explicit the different terms of the covariance, show how they are parametrised and how these parameters are optimised. We will then explain how this model can be used to assess the proportion of variance explained by each effect, as well as the statistical significance of each effect. See also **Fig. S1**.

We will rely on the following nomenclature and notations:

#### Nomenclature

*Molecule of interest*: Individual molecule, typically a gene or protein, on which SVCA is fitted.

*Cell state*: Intrinsic characteristic of a cell. In this paper, we take the overall expression profile excluding the gene or protein of interest as a multidimensional and continuous measure of cell state. Other possibilities include classifying cells into cell types.

*cellular neighbourhood matrix*: Continuous measure of the molecular composition of a cell’s neighbouring cells, summarised by weighting the molecular profiles of all neighbouring cells with a squared exponential function of their distance to the focal cell.

*Intrinsic effect*: Effect of the cell state on the expression level of the molecule of interest.

*Cell-cell interaction effect*: Effect of cell-cell interactions on the expression level of the molecule of interest. These interactions may harvest signaling between cells but also cell-types cooccurrences for example.

*Environmental effect*: Effect of the cell’s position on the expression level of the molecule of interest. This effect harvests unmeasured variables from the microenvironment with an effect on gene expressions, such as local glucose or oxygen access.

*Spatial variance signature*: Concatenation of all variance estimates (intrinsic effect, environmental effect, cell-cell interaction effect and residual noise) across all molecule for a given image.

#### Notations

- *N* - number of cells in a given image
- *D* - number of molecules (eg genes or proteins) in a given image
- *Y* - Expression level of the molecule of interest in all cells Dimensions: *N* × 1
- *X* - Cell state matrix made of the entire expression profile of each cell minus the molecule of interest. Dimensions: *N* (*D* 1). The molecule of interest is removed from the cell state matrix to prevent any cell-cell interaction false positive due to signal spillover between cells, as well as trivial intrinsic effect.
- *d_i, j_* - euclidean distance between cell *i* and *j*
- *K_int_* - cell-cell covariance for the intrinsic effect. Dimensions: *N* × *N*
- *K_c-c_* - cell-cell covariance for the cell-cell interaction effect. Dimensions: *N* × *N*
- *K_env_* - cell-cell covariance for the environmental effect. Dimensions: *N* × *N*

With these notations, SVCA models the expression level *Y* across cells as shown on Equation 1

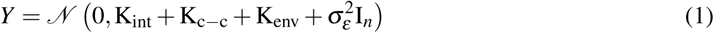

### 1.2 Definition of covariance terms

**Intrinsic effect:** The intrinsic effect is the effect of the cell state on the expression level of the molecule of interest. In our framework, it is modelled with the linear covariance term shown on Equation 2 and it resembles the Kinship matrix in GWAS studies.

The matrix X describes the cell states. If cell types are available, each row of X *X_n,_*: for cell *n* may be an indicator variable for the cell type of cell *n*. However, it is often preferable to not rely on uncertain and binary cell types and use a richer measure of cell states. In this paper, we use the expression profile of the cell, excluding the molecule of interest. This enables to account for any intrinsic correlation between genes.

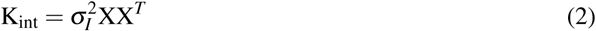

**Cell-cell interaction effect:** The cell-cell interaction effect harvests the effects of the types or states of all neighbouring cells on the expression level of the molecule of interest. In the GP framework, it is modelled with the covariance term shown on Equation 3.

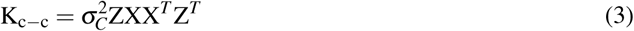

where

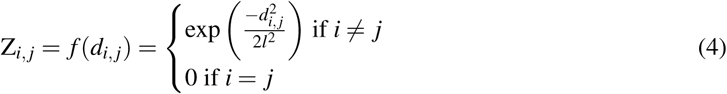

This covariance term is equivalent to a random effect model where, gene expression profiles of all neighbouring cells are used as predictor variables and the effect of a cell *i* on a cell *j* is weighted by a function of the distance between them: *Z_i, j_*, shown on equation 4.

**Environmental effect:** The environmental effect aims at harvesting other local sources of variation in the cell micro-environment which are not measured in the data and have an effect of the expression level of the modelled gene. To model this unobserved source of variation, we use a Squared Exponential Kernel (Eq 5) for its non-linearity and its successful application to spatial expression data. [23]

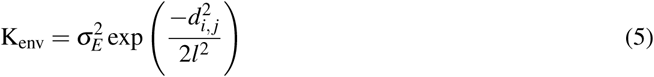

### 1.3 Parameters’ inference

The variance parameters of the SVCA model are optimised by maximising the log likelihood in Equation 6 [16, 13]. The scales of the covariance terms, *s_I_, s_C_, s_E_* and *s_e_* are optimised with gradient descent using a lbfgs optimiser, which updates the parameters iteratively through small steps along the gradient of the likelihood until it reaches a local optimum (null gradient). The length scale of the environmental and the local terms is optimised with a grid search strategy, to avoid possible local optima.

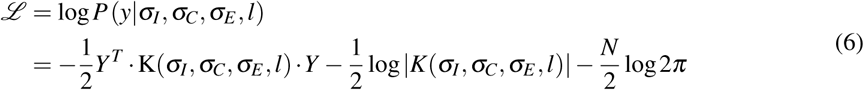

with

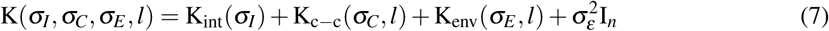

### 1.4 Estimates of variance components

Variance components for each effect are estimated using Gower factors[19, 12] (Equation 8).

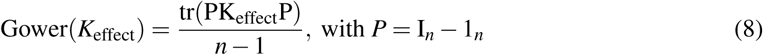

The Gower Factor of a covariance term computes the expected variance of a random variable which is normally distributed with the considered covariance (Equation 9). In other words, the Gower factor of each covariance term of the SVCA model computes the amount of gene/protein variance across cells which is explained by the corresponding effect.

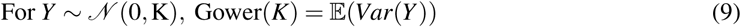

To compute the fraction of variance explained by each effect modelled in SVCA (intrinsic, environmental, cell-cell interactions and noise), we normalise Gower factors as shown on Equation 10

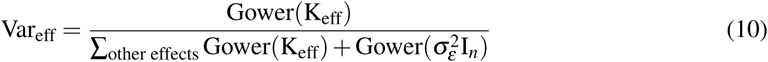

This procedure enables us to break down the variance of every protein, across cells in the three effects of interest plus the noise.

### 1.5 Significance of variance components

In this section, we explain how we assessed the significance of the cell-cell interaction component, as this is the variance component of main interest for our study. The significance of other variance components can be tested analogously.

#### 1.5.1 General Approach

We assessed the significance of the cell-cell interaction component using a log likelihood ratio (LLR) between the SVCA model and a reduced model omitting the cell-cell interaction component (Equation 11). Given that the reduced model is nested in the full SVCA model, we relied on Wilks’ theorem [24], which states that if the null hypothesis is true (no cell-cell interactions), the LLR statistics should follow a *c*2 distribution. In practise, we calibrate this *c*2 distribution by fitting its parameter to an empirical null distribution of LLRs obtained from simulations [5, 7].

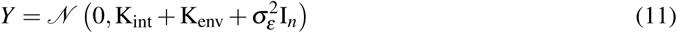

#### 1.5.2 Estimation of the null LLRs distributions

The simulation procedure is as follows. For all proteins and all images, we fitted the null model (Equation 11), and simulated data from the fitted normal distribution. We simulated 100 data points for each test and then fitted a *c*2 distribution to those using an off-the-shelf non linear optimisation method [11]. We then compared the LLR obtained, for each protein and each image, for the real data to the corresponding fitted *c*2 distribution and estimated p-values from this comparison.

#### 1.5.3 Multiple testing correction and False Discovery Rate

For every test, we computed a cell-cell interaction p value using the method described before and used the Benjamini-Hochberg procedure [4] to adjust p values for multiple testing. For each protein, we then counted the number of images in which the cell-cell interaction component was significant for a FDR threshold of 1%.

### 1.6 Comparison to related models

In this section, we describe the relationship between SVCA and existing modelling frameworks to quantify cell-cell interactions in spatially resolved single cell molecular data.

#### 1.6.1 Schapiro et al 2017 - HistoCAT

HistoCAT [17] aims at measuring spatial cooccurrence of different cell types. Briefly, cells of one or multiple images are classified into discrete cell-types based on their expression profile using a clustering algorithm. For every cell, a neighbourhood is defined as containing all cells within a fixed distance threshold (measured from membrane to membrane). Using this fixed neighbourhood definition, histoCAT counts the number of occurrences of a given pair of cell types, in the same neighbourhood. This number is then compared to a null distribution obtained from permuting the cells’ positions, which gives a p-value for positive and negative cell types interactions.

Unlike SVCA, histoCAT does not quantify the effect of these interactions on individual expression levels.

#### 1.6.2 Battich et al 2015

Battich et al [3] uses a regression approach to measure the effect of the cell microenvironment on individual expression levels. Briefly, 183 features are collected, quantifying intrinsic cell properties and microenvironmental properties. Microenvironmental features namely account for local cell crowding, number of adjacent neighbours, intercellular space around the cell, as well as the molecular profile of the neighbours, based on a fixed distance threshold. The dimensionality of this feature set is then reduced using principal component analysis (PCA), and single cell expression profiles are modelled with a fixed effect linear model with the first 20 PCs as covariates. The PCs are then a posteriori linked to the microenvironmental features of interest. Biological replicates are used to quantify the amount of variance explained by each covariate using out of sample prediction.

This method therefore quantifies directly the effect of microenvironmental features including cell-cell interactions. Unlike SVCA however, it relies on a definition of discrete microenvironmental features and the definition of fixed parameters such as a distance threshold to define a cell’s neighbourhood, which limits the applicability of the method to general spatial data.

#### 1.6.3 Goltsev et al 2018

Goltsev et al’s [1] approach also relies on the definition of discrete microenvironmental variables, used in a fixed effect linear model to predict the expression level of individual markers out of sample. In contrast to Battich et al [3], microenvironmental variables are not defined directly based on the molecular profile of neighbouring cells, but based on the cell-type composition of the neighbourhood. The different neighbourhood cell-type compositions are clustered into discrete *i-niches*, used as a discrete input for the linear model.

This method therefore enables to quantify directly the effect of cell-cell interactions on individual molecular profiles of single cells. However, it again relies on a priori definition of microenvironmental variables, this time based on discrete cell-type assignments.

## 2 Model Validation

### 2.1 Simulations of the cell-cell interaction component

In order to be as realistic as possible, our simulations were based on real data from 11 images of the IMC dataset [10]: real cell positions, cell states, intrinsic and environmental effects were used, and only the cell-cell interaction effect was rescaled for the purpose of the simulations.

Our workflow was as followed:

1. Fitting the SVCA model to the real dataset considered here (11 images and 26 proteins)
2. Simulating data from a multivariate normal distribution, with a covariance made of:

- the intrinsic covariance from the fitted model
- the environmental covariance from the fitted model
- the noise covariance from the fitted model
- a cell-cell interaction covariance which is a **rescaled version** of the one fitted to the data (Equation 12)
3. Reffiting SVCA to the simulated data, where the variance explained by cell-cell interactions is known from the rescaling step.
4. Comparing the variance estimates for cell-cell interactions with the ground truth.

In Equation 12, the proportion of variance attributable to cell-cell interactions in the simulated data, *x* ranged from 10% to 90% (*x* ϵ [0.1; 0.9]), and the rescaling factor *k_sim_*was chosen accordingly.

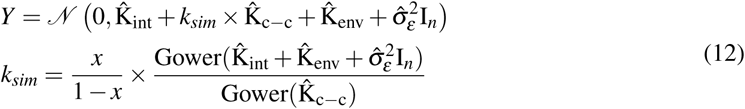

### 2.2 P-value calibration for the cell-cell interaction component

We used simulations to assess the p-value calibration for the cell-cell interaction component. For 11 random images and all 26 proteins of the IMC dataset, we simulated from the null model (SVCA without the cell-cell interaction term whose parameters were fitted to the data). We then used the procedure described above to compute p-values for the null distribution, and computed the empirical false positive rate for multiple p-value thresholds (**Fig. 2**).

### 2.3 Analysis of model robustness

We used a bootstrapping strategy to quantify the uncertainty on the variance estimates. For a given image and a given protein, the model is fitted 5 times on a different subset of 80% of the cells. In order to visualise the robustness of SVCA signatures for all images at once, we then represented signatures computed from each bootstrap and each image in a low dimensional latent space using t-SNE in order to compare visually the distance between signatures coming from multiple bootstraps of the same image and signatures coming from different images (**Fig. S4-S12-S15**).

#### 2.4 Out of sample prediction

We used 5-fold cross-validation to assess predictions of the expression profiles using alternative models. In order to check whether every covariance term increased the predictive power of the model, we considered the following variance component models:

1. a model with only an intrinsic covariance to it.
2. a model with an intrinsic component and a local component.
3. the full model with all three terms

### 2.5 Identifiability of cell-cell interactions versus environmental effects

To better understand the identifiability of cell-cell interactions versus environmental effects, we compared the variance estimates of SVCA with the variance estimates of a reduced model which does not account for cell-cell interactions (Equation 11). Both models were fitted in the simulation setting described in the main text (26 proteins, 11 images and 10 repeat experiments). Variance estimates of SVCA and the reduced model were averaged across all proteins, images and experiments.

We then built a Sankey plot (**Fig. 2e**) to visualise what terms of the reduced model captured the residual variance from the lack of cell-cell interaction modelling. The width of the edges correspond to the increase in the variance estimates for the intrinsic effect, the environmental effect and the noise, from the SVCA model to the reduced model. This represents the redistribution of the cell-cell interaction component to other variance estimate from the SVCA model to the reduced model.

## 3 Downstream Analysis

### 3.1 Variance across images of the IMC dataset and patient effect

We also asked how the variability of the results across the IMC images related to the patient the biopsy came from (so called patient effect, **Fig. S3**). In order to do that, we fitted the following variance decomposition model for each protein and each variance component:

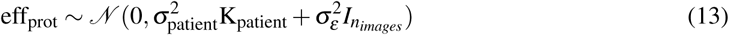

where

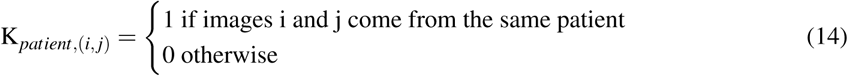

The percentage of variance explained by the patient effect was then computed with Gower factors.

### 3.2 Gene Set Enrichment Analysis

When the number of genes allowed (seq-FISH), we ran a gene set enrichment analysis based on the ranking of the cell-cell interaction component of the spatial variance signature. The aim here was to understand the types of biological processes which are influenced by cell-cell interactions.

A simple procedure based on the idea of GSEA [22, 15] was implemented and used:

1. Genes are ranked based on the size of the cell-cell interaction component in the Spatial Variance signature
2. Gene sets are taken from the reactome database [8, 9] and are filtered for pathways including at least five genes of the gene set analysed in the seq-FISH data
3. A GSEA-like trace is computed for each pathway and the height of this trace is considered as a test statistic.
4. Gene names are permuted 10,000 times in order to estimate an empirical p-value for the statistic described above.
5. p-values are adjusted for multiple testing using a Benjamini-Hochberg procedure [4]

## 4 Dataset

### 4.1 Dataset description

#### 4.1.1 Imaging Mass Cytometry (IMC) data

With IMC, the analysed tissue or cell culture is laser-disected into a sub cellular resolution grid of so called voxels of dimension 1*µm* 1*µm*. Every voxel of this grid is then analysed with cyTOF (anti-body based method), which results in protein counts of 26 proteins per voxel, which can be aggregated into single cell counts after cell segmentation [10, 21, 18, 6].

We analysed a dataset of 52 breast cancer biopsies imaged with Imaging Mass Cytometry coming from 27 patients [17]. 41 of these images are associated to clinical data:

*•* ER status
*•* PR status
*•* Her2 status
*•* Grade
*•* Biopsy location (periphery or centre)

These images contain between 267 and 1455 cells, with an average around 900 cells. 26 proteins counts are quantified at a sub cellular level (between 10 and 100 pixels/measurements per cell). These proteins are:

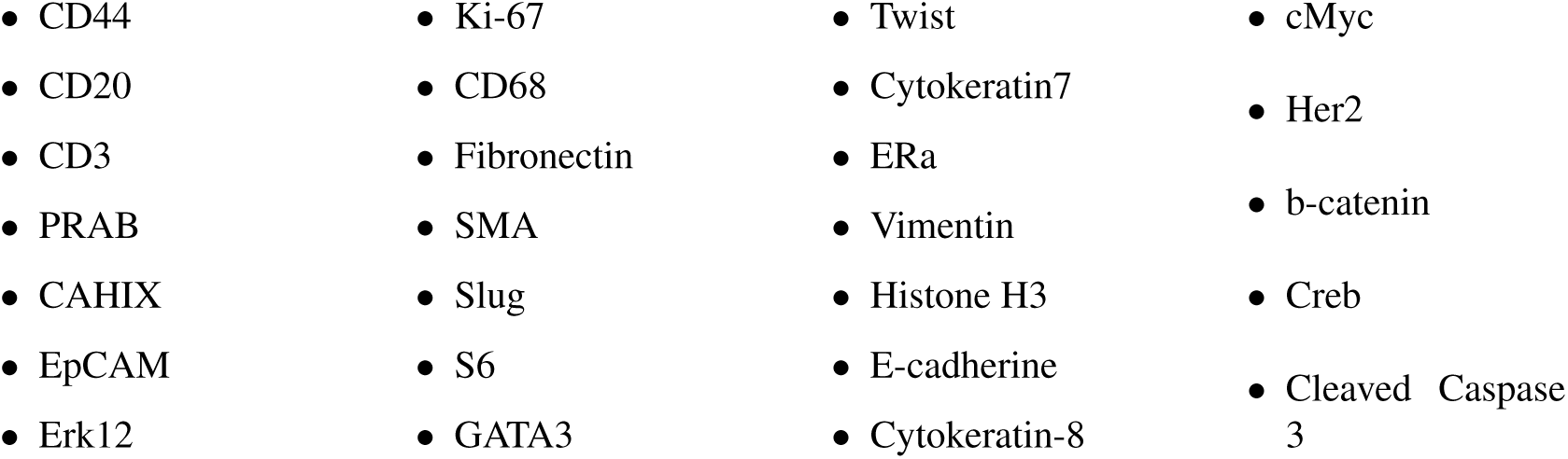

#### 4.1.2 mer-FISH and seq-FISH data

Although mer-FISH and seq-FISH techniques differ slightly, the data produced and available online [14, 20] come in a similar format. Briefly, it comes as a list of detected individual RNA molecules, associated to a precise position on the tissue and the index of the cell each molecule belongs to (obtained with automatic cell segmentation). Summarising this data into a molecule count at the single cell level is therefore straightforward.

We analysed a mer-FISH dataset of 20 images taken on a single plate of breast cancer cell culture. Each image contained between 2500 and 2900 cells and 130 genes were measured. Additionally, we analysed a seqFISH dataset consisting of 20 images of a single mouse hippocampus [20]. The images were taken in different regions of the hippocampus and 249 genes were measured.

### 4.2 Data processing

For the IMC dataset, single cell expression levels were computed by taking the median protein count across pixels. The median was chosen instead of the mean or the sum for its greater robustness against technical outliers produced by the experimental pipeline. For the mer-FISH and the seq-FISH datasets, single cell RNA counts were obtained directly from summarising the data described before.

In all cases, the data was then transformed with an Anscombe’s transformation for variance stabilisation of Negative Binomial data [2]. The dispersion parameter *f* in Equation 15 is optimised with gradient descent and the log transformation of Equation 16 is applied to the data.

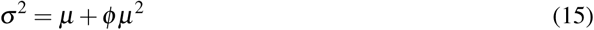

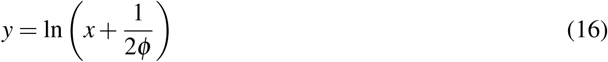

The resulting signal is then normalised by regressing out the log of the total signal in the cell. This last step aims at taking into account local batch effects which would make some cells “brighter” overall.

Before fitting the SVCA model on a given protein or gene, the stabilised expression levels are subsequently raked standardised and transformed into normally distributed data using the probit function. This ensures a more robust fitting process due to a lesser sensitivity to outliers.

